# Gain and loss of gene function shaped the nickel hyperaccumulation trait in *Noccaea caerulescens*

**DOI:** 10.1101/2025.07.22.664359

**Authors:** Célestine Belloeil, Vanesa Sanchez Garcia de la Torre, Rubén Contreras-Aguilera, Hendrik Küpper, Céline Roques, Carole Iampietro, Céline Vandecasteele, Christophe Klopp, Alexandra Launay-Avon, Wiebke Leemhuis, Jitpanu Yamjabok, Joost van den Heuvel, Mark G.M. Aarts, Celestino Quintela-Sabarís, Sébastien Thomine, Sylvain Merlot

## Abstract

Nickel hyperaccumulation is an extreme adaptation to ultramafic soils observed in more than 500 plant species. However, our understanding of the molecular mechanisms underlying the evolution of this trait remains limited. To shed light on these mechanisms, we have generated a high-quality genome assembly of the metal hyperaccumulator *Noccaea caerulescens*. We then used this genome as reference to conduct comparative intraspecific and interspecific transcriptomic analyses using various accessions of *N. caerulescens* and the non-accumulating relative *Microthlaspi perfoliatum*, to identify genes associated with nickel hyperaccumulation. Our results suggest a correlation between nickel hyperaccumulation and a decrease in the expression of genes involved in defense responses and the regulation of membrane trafficking. Surprisingly, these analyses did not reveal a significant enrichment of genes involved in the regulation of metal homeostasis. However, we found that the expression levels of selected metal transporters, namely NcHMA3, NcHMA4 and NcIREG2, is consistently elevated in *N. caerulescens* accessions hyperaccumulating nickel. Furthermore, our analyses identified frameshift mutations in NcIRT1 associated with the loss of nickel hyperaccumulation in a few accessions. We further showed that the expression of a functional NcIRT1 in roots of the La Calamine accession increases nickel accumulation in shoots. Our results demonstrate that NcIRT1 participate in nickel hyperaccumulation in *N. caerulescens.* They also suggest that nickel hyperaccumulation is an ancient trait in *N. caerulescens* that has evolved from the high and constitutive expression of few metal transporters including *NcIREG2* and that the trait was subsequently lost in a few accessions due to mutations in NcIRT1.

## Introduction

The ability of plant species to accumulate very high amounts of metals has piqued the interest of biologists since the discovery of the first metal hyperaccumulator species more than 150 years ago (Sachs, 1865; Minguzzi C and Vergnano O, 1948; Jaffré et al., 1976; Severne and Brooks, 1972). Today, in the context of the energy transition greedy for metals, these plants represent an opportunity to develop sustainable technologies aimed at remediating metal-contaminated soils, thereby helping to mitigate the environmental impacts of the metal industry (van der Ent et al., 2015; Suman et al., 2018; Rylott and van der Ent, 2025).

The majority of plant species that are adapted to toxic metalliferous soils, of geological or anthropogenic origin, exclude metals from their photosynthetic tissues. Only a few species, known as hyperaccumulators, have evolved the ability to accumulate metals in leaves up to 100 to 1000 times the concentrations measured in other plant species, without showing signs of toxicity (Baker, 1981; van der Ent et al., 2013). This phenomenon is regarded as an extreme adaptation to metalliferous soils, providing selective advantages such as allelopathy or elemental defense against pathogens and insect herbivory (Manara et al., 2020). To date, over 700 hyperaccumulator species have been described that accumulate nickel, manganese, zinc, or cadmium. However, nickel hyperaccumulators, comprising more than 500 species across over 50 families, represent the vast majority and diversity of these peculiar species (Reeves et al., 2018). The prevalence of nickel hyperaccumulators can be explained by the relative abundance of large areas of nickel-rich ultramafic soils, including serpentine outcrops, on the surface of the Earth compared to the more restricted area of exposed zinc- and cadmium-rich calamine soils (Krämer, 2010; Garnica-Díaz et al., 2023; Baker et al., 2010).

Despite the predominance of nickel hyperaccumulators, most of our knowledge on the mechanisms of metal hyperaccumulation comes from the study of zinc and cadmium hyperaccumulation in the Brassicaceae species *Arabidopsis halleri* and *Noccaea caerulescens* (formerly known as *Thlaspi caerulescens*). It has been proposed that the evolution of metal hyperaccumulation resulted from the enhanced activity of genes normally involved in the acquisition and homeostasis of metals, essentially in three determinant steps: the uptake of metals by roots, the translocation of metals from roots to shoots, and the detoxification and sequestration of metals in shoots (Leitenmaier and Küpper, 2013; Manara et al., 2020; Krämer, 2010; Verbruggen et al., 2009). Accordingly, comparative transcriptomic analysis has revealed that zinc and cadmium hyperaccumulation is associated with the high and constitutive expression of genes involved in metal transport, including heavy metal ATPases (HMA), ZRT/IRT-like protein (ZIP), cation diffusion facilitators/metal tolerance proteins (CDF/MTP), and in the synthesis of metal chelators such as nicotianamine synthases (NAS) (van de Mortel et al., 2006; Hammond et al., 2006; Becher et al., 2004; Talke et al., 2006). Furthermore, the role of orthologs of HMA3, HMA4, and NAS2 in the hyperaccumulation of zinc and cadmium was confirmed by the silencing of the corresponding genes in *A. halleri* and in the Crassulaceae species *Sedum plumbizincicola* (Hanikenne et al., 2008; Deinlein et al., 2012; Liu et al., 2017).

More recently, comparative transcriptomic studies have been performed in species from different plant families to identify genes associated with nickel hyperaccumulation (Meier et al., 2018; Halimaa et al., 2014b; García de la Torre et al., 2021). These studies revealed several candidate genes involved in metal transport, responses to biotic and abiotic stresses, biosynthesis of specialized metabolites, or DNA modifications. In particular, the high expression of genes encoding metal transporters of the ZIP and the Iron Regulated/Ferroportin (IREG/FPN) families, was repeatedly associated with nickel hyperaccumulation in different plant families. Furthermore, the silencing of *NcIREG2* in roots of *N. caerulescens* provided direct genetic evidence that this transporter is important for nickel transport and accumulation in vacuoles (García de la Torre et al., 2021).

*Noccaea caerulescens* has been proposed for several years as an alternative genetic model to study the mechanisms of metal hyperaccumulation and their evolution, because in contrast to *A. halleri*, which is able to hyperaccumulate zinc and cadmium, *N. caerulescens* accessions are also able to hyperaccumulate nickel (Assunção et al., 2003b; Peer et al., 2003; Hanikenne and Nouet, 2011).

*N. caerulescens* is a diploid, self-fertile species of the Brassicaceae tribe Coluteocarpeae. This tribe also include the genus *Microthlaspi* that diverged from *Noccaea* about 6 Ma ago (Hendriks et al., 2023). *N. caerulescens* is commonly found on non-metalliferous and on the metalliferous calamine and ultramafic soils throughout Europe, where it consequently hyperaccumulates zinc, cadmium or nickel (Jakovljević et al., 2024; Assunção et al., 2003a). In contrast, the *Microthlaspi* species, *M. erraticum* and *M. perfoliatum*, develop mainly on limestone and do not accumulate metals (Mishra et al., 2020; Ali et al., 2019; Kozhevnikova et al., 2020). When grown *ex situ*, *N. caerulescens* accessions show variable capacities to tolerate and accumulate metals (Seregin et al., 2022; Kozhevnikova et al., 2020; Sterckeman et al., 2017; Assunção et al., 2003a). However, while zinc hyperaccumulation appears to be a species-wide trait in this species, the ability to hyperaccumulate nickel is absent in a few accessions (*e.g.* La Calamine, Prayon, Plombières) developing on calamine soils in Belgium (Kozhevnikova et al., 2020; Seregin et al., 2022). A recent population genetic study revealed that *N. caerulescens* accessions found in western Europe belong to at least three distinct genetic units originating from two populations that diverged during the last glacial period in the Apennine and the Iberian refugia. The later population further diverged into a third genetic unit during the recolonization of northern Europe (Gonneau et al., 2017). Accessions adapted to ultramafic soils and hyperaccumulating nickel (e.g. Puente Basadre, Bergenbach and Firmiensis, also known as Puy de Wolf) are found in all these three genetic units. This suggests that the adaptation to ultramafic soils and the associated nickel hyperaccumulation trait has either evolved repeatedly in the different genetic units, or is an ancestral trait in *N. caerulescens* that was subsequently lost in some accessions.

The identification of the mechanisms involved in nickel hyperaccumulation and the study of their evolution in *N. caerulescens* require a better knowledge of the *N. caerulescens* genome. This genome is derived from the proto-Calepineae karyotype, and is composed of seven chromosomes with an estimated size of 270 Mb (Mandáková et al., 2015). However, the only genome sequence publicly available for this species was assembled using short-reads, resulting in more than 19,000 scaffolds with a total size of 140 Mb, indicating that this assembly is still highly fragmented and incomplete (Kiefer et al., 2019).

In this work, we produced a high-quality assembly of the *N. caerulescens* genome, using long read sequencing technology, which we subsequently used as a reference to identify differentially expressed gene and genetic polymorphisms associated with the capacity to hyperaccumulate nickel in the Coluteocarpeae tribe. Our results indicate that nickel hyperaccumulation in *N. caerulescens* is associated with the increased expression of a few genes encoding metal transporters, while the absence of the trait in few accessions is linked to the loss of function of the NcIRT1 transporter. We further provide functional and genetic evidence for a role of NcIRT1 in nickel uptake in *N. caerulescens*. Our results suggest that nickel hyperaccumulation is an ancestral character in *N. caerulescens* that likely co-evolved with the zinc hyperaccumulation trait.

## Results

### A chromosome-level assembly of the Noccaea caerulescens genome

To enable the dissection of the molecular mechanisms underlying nickel hyperaccumulation in *N. caerulescens*, we undertook the genome sequencing of the Cira accession (*Ncci*), originating from ultramafic outcrops in Galicia, Spain. We sequenced this genome using the Pacbio Hifi technology (Table 1). The k-mer analysis of the Hifi reads predicted a genome size of 303 Mb with 61% of repeated sequences. The assembly generated 1206 contigs with a total size of 295.6 Mb (97.2% of the theoretical genome size). In this initial assembly, the 23 largest contigs, ranging from 3 to 33 Mb, covered 80% of the genome assembly. To further improve the assembly of the *Ncci* genome, we ordered and oriented the contigs in chromosomes using Hi-C chromatin conformation capture reads. This scaffolding step generated seven large scaffolds, representing the seven chromosomes of the *N. caerulescens* genome, covering 85% of the total assembly (Fig. 1). The annotation of this genome assembly (v2.3) predicted 31794 protein-coding genes, of which 82% and 71% were annotated by Gene Ontology (GO) and Mercator4 (Supplementary Data Set S1). The seven large scaffolds contain 96% of the predicted genes, indicating that non-scaffolded contigs are mainly composed of highly repetitive non-coding sequences. The examination of the functional annotation of the *Ncci* genome assembly revealed the duplication of several genes involved in solute transport and nutrient uptake, including orthologs of *HMA3* (g5553 and g29551), *HMA4* (g10683 and g10685), *NRAMP1* (g8380 and g8418)*, IRT2* (g27496, g27497 and g27498) and *ZIP12* (g19758, g19882, g19892 and g19949), which may reflect the adaptation of *N. caerulescens* to metalliferous soils (Fig. 1; Supplementary Data Set S1). This first high-quality annotated genome assembly of *N. caerulescens* provides a robust basis for subsequent transcriptomic and genomic intra- and interspecific comparisons.

**Table 1.** Descriptive statistics of the *Noccaea caerulescens* Cira genome assembly and annotation (v2.3).

|  |  |
| --- | --- |
| Assembly category | *N. caerulea* Cira v2.3 |
| **k-mer analysis** |  |
| Genome haploid length (k = 21) | 303.9 Mb |
| Genome repeat length | 186.5 Mb (61 %) |
| **Hifi read assembly** |  |
| Number of contigs | 1206 |
| Total size contigs | 295.6 Mb |
| Longest contig | 33.1 Mb |
| N50 contig length | 12.1 Mb |
| L80 contig number | 23 |
| **Hi-C scaffolding** |  |
| Number of scaffolds | 1175 |
| Longest scaffold | 46 Mb |
| Number of scaffolds > 10 Mbp | 7 |
| N50 scaffold length | 36.2 Mb |
| L85 scaffold number | 7 |
| **Annotation** |  |
| BUSCO score | C: 97.3 % [S: 91.9 %, D: 5.4 %] |
| Number of genes | 31 794 |
| Number of genes in the 7 largest scaffolds | 30 667 (96 %) |
| Number of annotated genes (GO) | 26 142 (82 %) |
| Transposable elements | 57 % |

GO : Gene Ontology

**Figure 1.**
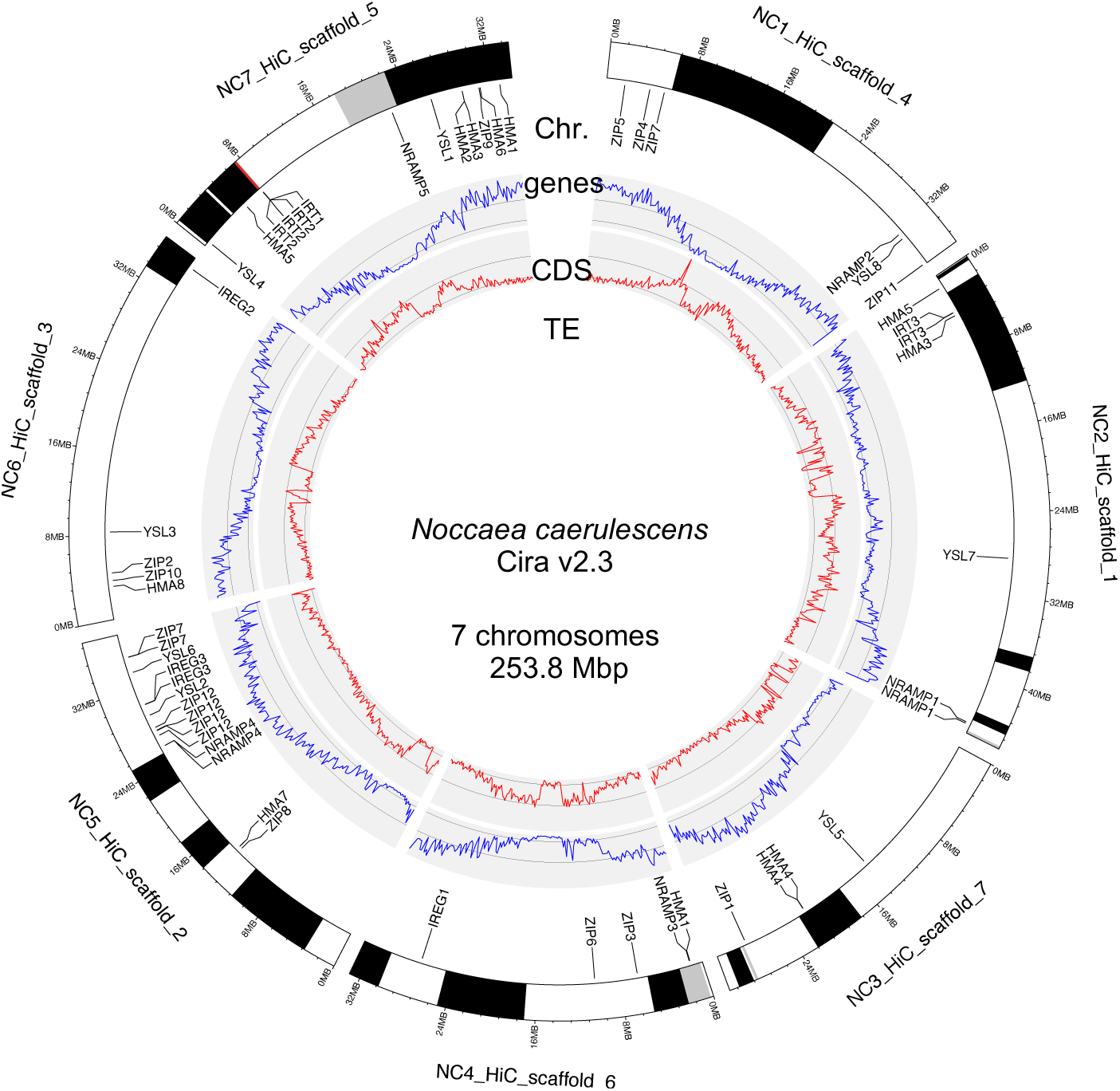
Graphical representation of the *Noccaea caerulescens* Cira genome assembly (v2.3). The outermost ring represents the largest scaffolds composed of Hifi contigs (colored bands), and representing the seven chromosomes of the *N. caerulescens* genome. The chromosomes (Chr.) are annotated with the position of genes encoding for metal transporters of the ZIP/IRT, HMA, NRAMP, YSL and IREG families (genes). Inner rings successively represent the density distribution of coding sequence (CDS) and transposable elements (TE) using a 200 kb window.

### Transcriptomic response to nickel in *N. caerulescens* and *M. perfoliatum*

To identify putative mechanisms involved in nickel tolerance and accumulation in *N. caerulescens*, we first searched for differentially expressed (DE) genes in response to nickel in the Firmiensis accession (*Ncfi*), also known as Puy de Wolf, adapted to ultramafic soil and hyperaccumulating nickel. For nickel treatment, we chose a concentration of 37.5 µM, which is sufficient to reveal the hyperaccumulation trait of *Ncfi*, but does not visibly affect the growth of *M. perfoliatum* under our hydroponic growth conditions. Differential expression analysis revealed 4 and 7 DE genes (|log2 Fold Change (FC)| ≥ 1, FDR-adjusted p-value < 0.05, n=3) responsive to nickel in roots and shoots of *Ncfi*, respectively (Fig. 2, A and B; Supplementary Data Set S2). Functional annotation indicated that some of these genes may play a role in pathogen and stress responses [*e.g.* PR-4 (g11320), SAG12 (g16833), DTX47(g30574)], but none of them seemed to play a direct role in the regulation of metal transport and homeostasis. As a comparison, we analyzed the nickel response in *M. perfoliatum* adapted to calcareous soils. Since we calculated that *N. caerulescens* and *M. perfoliatum* share 94% sequence identity in coding regions, we could use the *N. caerulescens* genome assembly v2.3 as a reference to analyze gene expression in *M. perfoliatum*. We identified 104 and 98 DE genes (|log2FC| ≥ 1, FDR-adjusted p-value < 0.05, n=3) responsive to nickel in roots and shoots of *M. perfoliatum*, respectively (Fig. 2, C and D; Supplementary Data Set S2). Among the genes up-regulated in response to nickel in roots (34 genes) and shoots (18 genes), we observed an enrichment of genes involved in iron homeostasis including orthologs of genes encoding ferric reduction oxidases (g20193, g21593), IMA/FEP peptide precursor genes (g4087, g13949), and the feruloyl-CoA 6′-hydroxylase 1 (g12319) involved in coumarin biosynthesis. These results indicate that nickel induces an iron deficiency response in *M. perfoliatum*, in agreement with what was observed in *A. thaliana* (Lešková et al., 2020). In comparison, *N. caerulescens* appears to be essentially insensitive to nickel treatment at the transcriptomic level. Hence, the mechanisms allowing nickel tolerance and hyperaccumulation in *N. caerulescens* are likely resulting from the high and constitutive expression of genes at the species level, rather than from the induction of a nickel tolerance and accumulation response upon metal treatment, as already reported for Zn and Cd hyperaccumulation in *A. halleri* (Krämer, 2010; Verbruggen et al., 2009; Hanikenne and Nouet, 2011).

**Figure 2.**
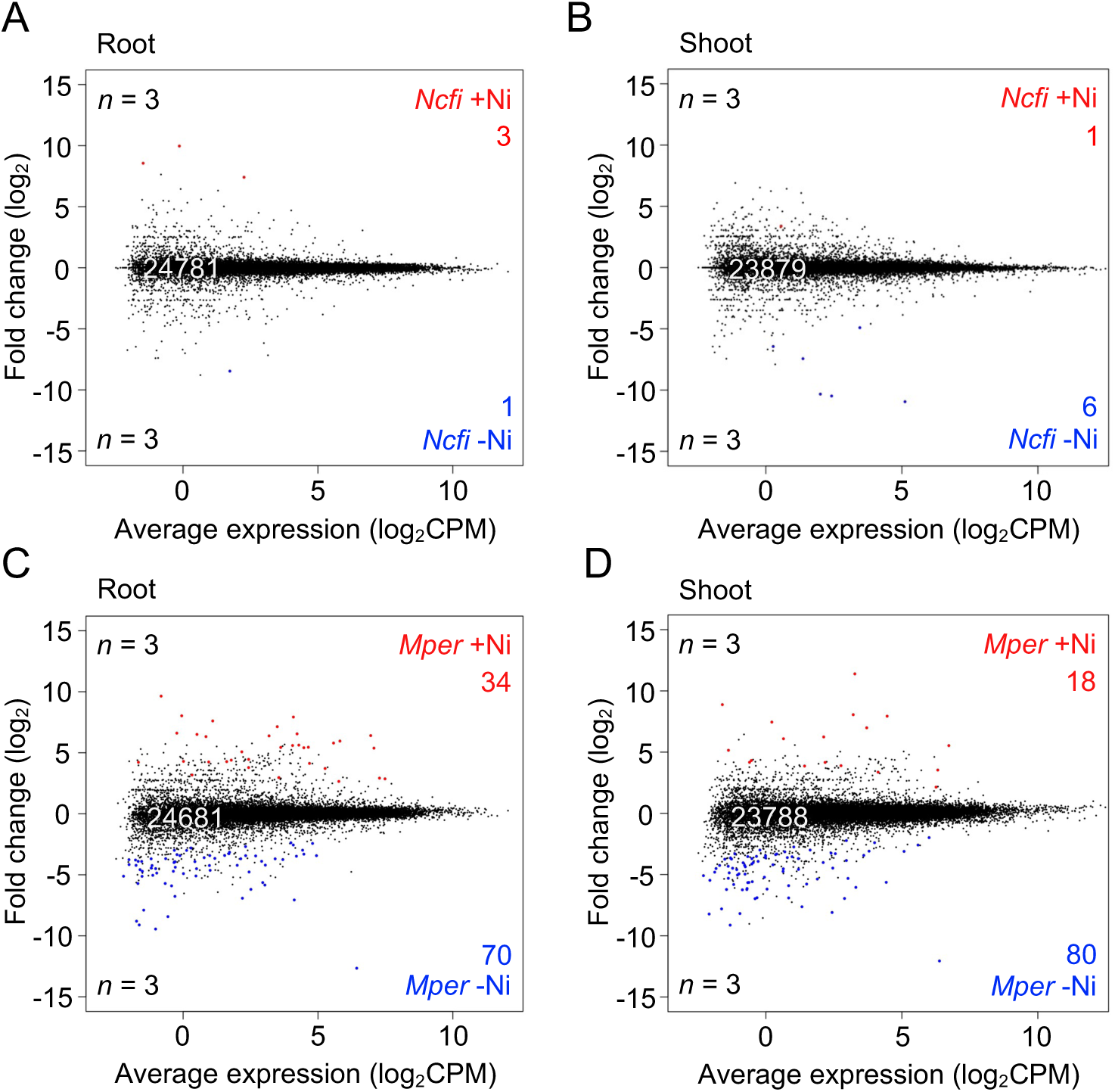
Identification of differentially expressed genes in response to nickel in *Noccaea caerulescens* and *Microthlaspi perfoliatum.* MA plots representing the comparative transcriptomic analysis of gene expressed in A) roots and B) shoots of *N. caerulescens* Firmiensis (*Ncfi*) in absence (-Ni) or presence of 37.5 µM nickel (+Ni). MA plots representing the comparative transcriptomics analysis of gene expressed in C) roots and D) shoots of *M. perfoliatum* (*Mper*) in absence or presence of nickel. In all comparisons, the number of differentially expressed genes (DEG) more expressed in response to nickel (red dots) or in absence of nickel (blue dots) are indicated (|log_2_FC|≥ 1, FDR adjusted p-value < 0.05, *n* = 3 biological replicates).

### Identification of genes associated with nickel hyperaccumulation in Noccaea caerulescens

To identify the constitutive transcriptomic differences accounting for nickel hyperaccumulation in *N. caerulescens*, we undertook an interspecific comparative transcriptomic approach. We sought to identify genes whose expression correlates with nickel hyperaccumulation using transcriptomic comparisons between three nickel hyperaccumulating accessions of *N. caerulescens* [i.e. Firmiensis (*Ncfi*), Bergenbach (*Ncbe*), and Puente Basadre (*Ncpu*)] and the non-accumulator species *M. perfoliatum* (Fig. 3; Supplementary Fig. S1; Supplementary Data Set S3). In each comparison, we identified between 7% and 10% of DE genes displaying a high fold change in the root and shoot transcriptomes (|log2FC| ≥ 3.32, FDR-adjusted p-value < 0.05, n=3). However, after combining the results of the three comparisons, we retained 855 and 931 common DE genes in roots and shoots, respectively (Fig. 3, C and D). Among the genes less expressed in the nickel hyperaccumulating accessions of *N. caerulescens*, we observed an enrichment for the functional categories corresponding to defense responses and membrane trafficking (Table 2), including the genes for guanine nucleotide-exchange protein BIG2 and the ADP-ribosylation related protein GB1 (Kitakura et al., 2017; Niu et al., 2022). We did not observe an enrichment of specific biological functions among the genes more highly expressed in *N. caerulescens*. Only a few genes encoding for orthologs of metal transporters are differentially expressed between nickel hyperaccumulating accessions of *N. caerulescens* and *M. perfoliatum*. The expression of the Heavy Metal ATPase *HMA4* genes (g10683, g10685) is 20 to 70 times higher in roots and 30 times higher in shoots of *N. caerulescens* compared to *M. perfoliatum*. In addition, the expression of *HMA3* genes (g5553, g29551) is between 25 and 200 times higher in shoots of *N. caerulescens*. The expression of the IREG/ferroportin transporter gene *NcIREG2* (g26066) is 30 times higher in *N. caerulescens* shoots compared to *M. perfoliatum*. Furthermore, read count analysis showed that the genes encoding these transporters are among the most highly expressed in *N. caerulescens* (Supplementary Data Set S3). These results indicate that the nickel hyperaccumulation trait is consistently associated with the high expression of only a few metal transporters in *N. caerulescens* when compared to the closely related non-accumulating species *M. perfoliatum.* Orthologs of HMA3 and HMA4 that have previously been identified as decisive for cadmium and zinc hyperaccumulation in both *A. halleri* and *N. caerulescens* are also constitutively highly expressed in nickel hyperaccumulating accession of *N. caerulescens* (Becher et al., 2004; Bernard et al., 2004; Papoyan and Kochian, 2004; Mishra et al., 2017). In contrast, the constitutive high expression of IREG2 orthologs in shoots is a hallmark of nickel hyperaccumulators, supporting the specific role of this transporter in nickel hyperaccumulation in *N. caerulescens* (Halimaa et al., 2014a; Meier et al., 2018; García de la Torre et al., 2021).

**Figure 3.**
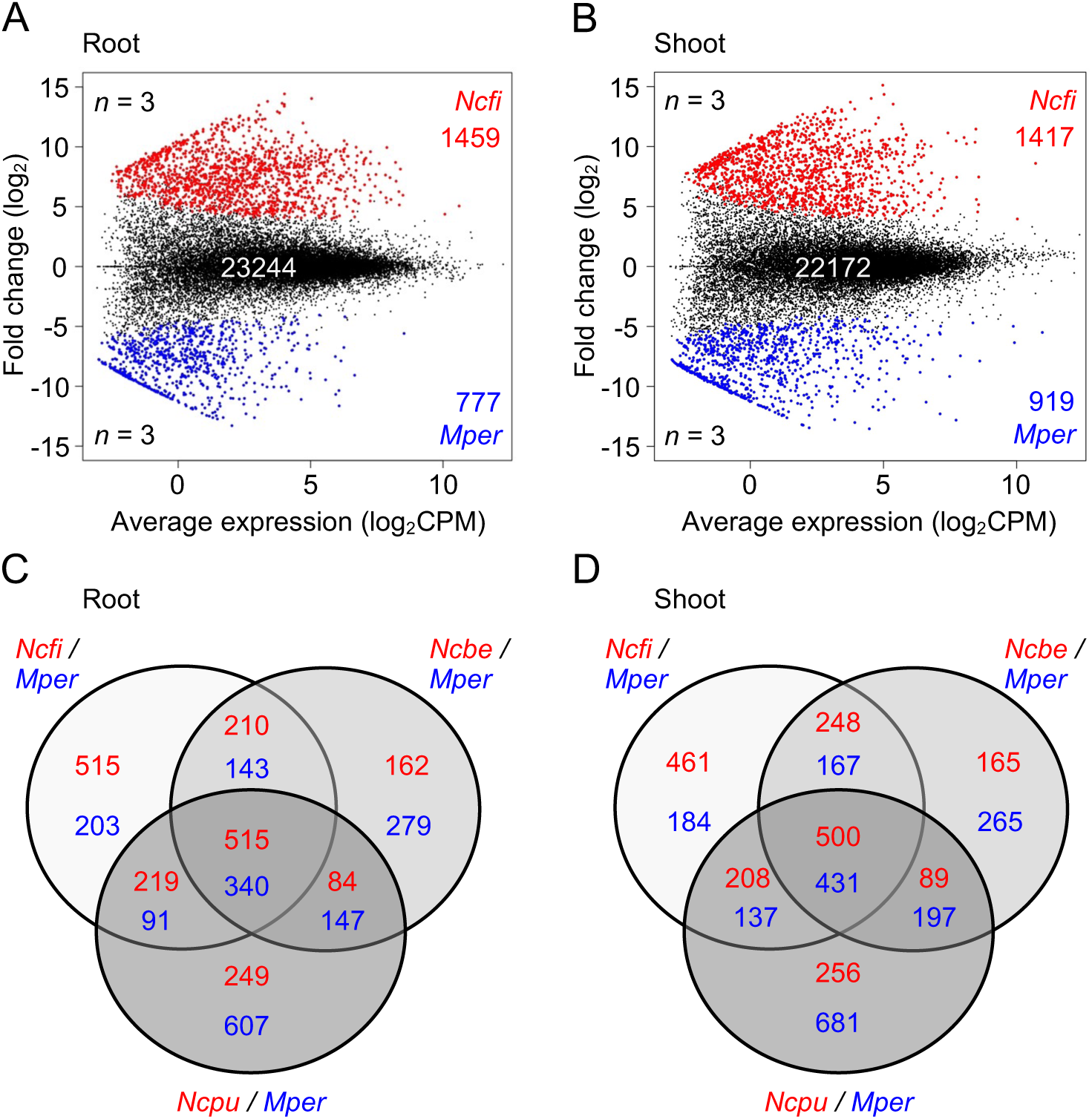
Identification of differentially expressed (DE) genes between *Noccaea caerulescens* and *Microthlaspi perfoliatum.* A) MA plot representing the interspecific comparative transcriptomic analysis of gene expressed in roots of *N. caerulescens* Firmiensis (*Ncfi*) and *M. perfoliatum* (*Mper*). B) MA plot representing the interspecific comparative transcriptomic analysis of gene expressed in shoots of *Ncfi* and *Mper*. In these comparisons, the number of DE genes (|log_2_FC|≥ 3.32, FDR adjusted p-value < 0.05, *n* = 3 biological replicates), more expressed in *Ncfi* (red dots) or in *Mper* (blue dots) are indicated. The comparisons of *N. caerulescens* accessions Bergenbach (*Ncbe*) and Puente Basadre (*Ncpu*) with *M. perfoliatum* are presented in Supplementary Fig. S1. C) Venn diagram displaying the number of DE genes in roots between *Ncfi*, *Ncbe* and *Ncpu* (red) and *Mper* (blue). D) Venn diagram displaying the number of DE genes in shoots between *Ncfi*, *Ncbe*, *Ncpu* (red) and *Mper* (blue).

**Table 2.**
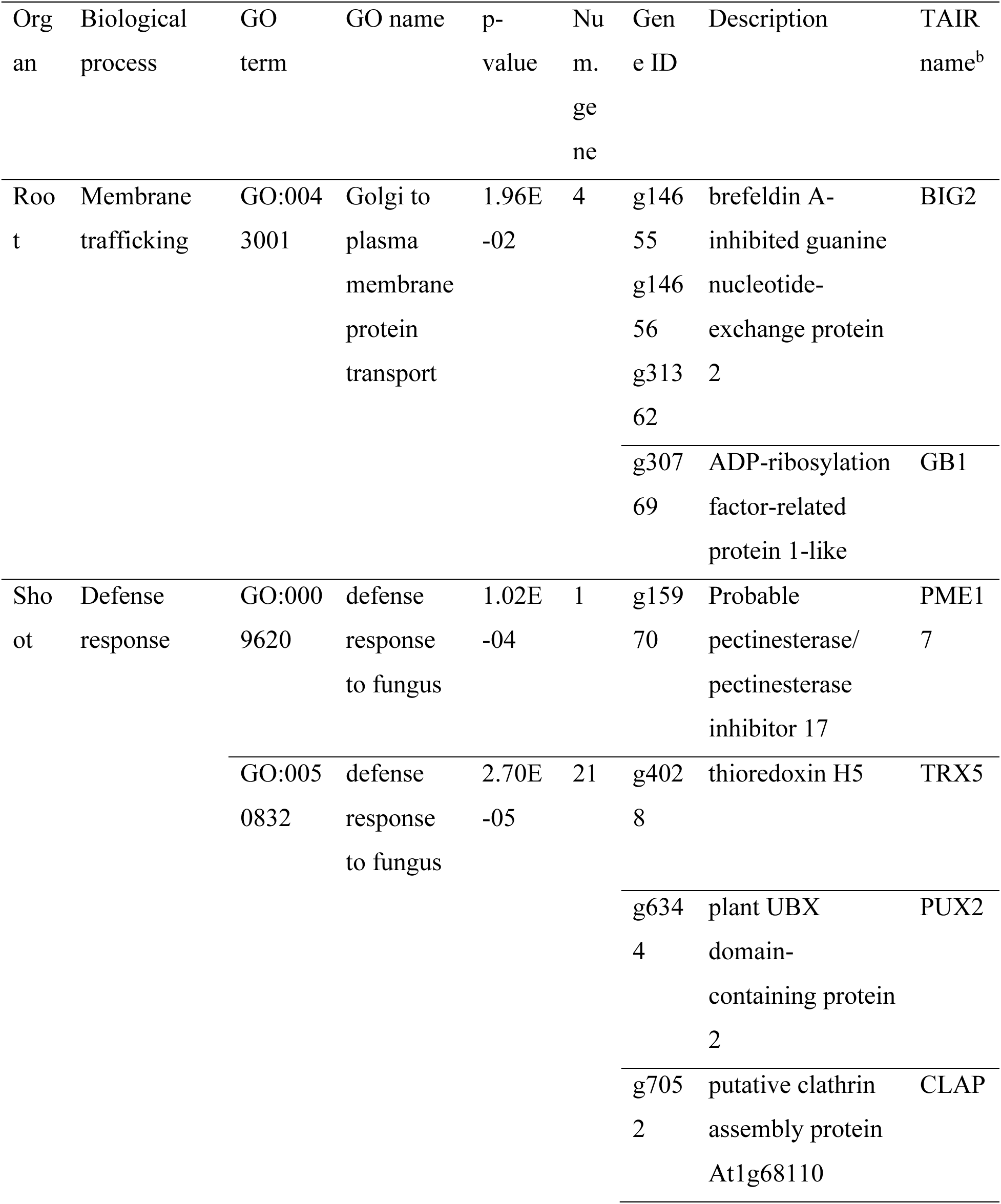
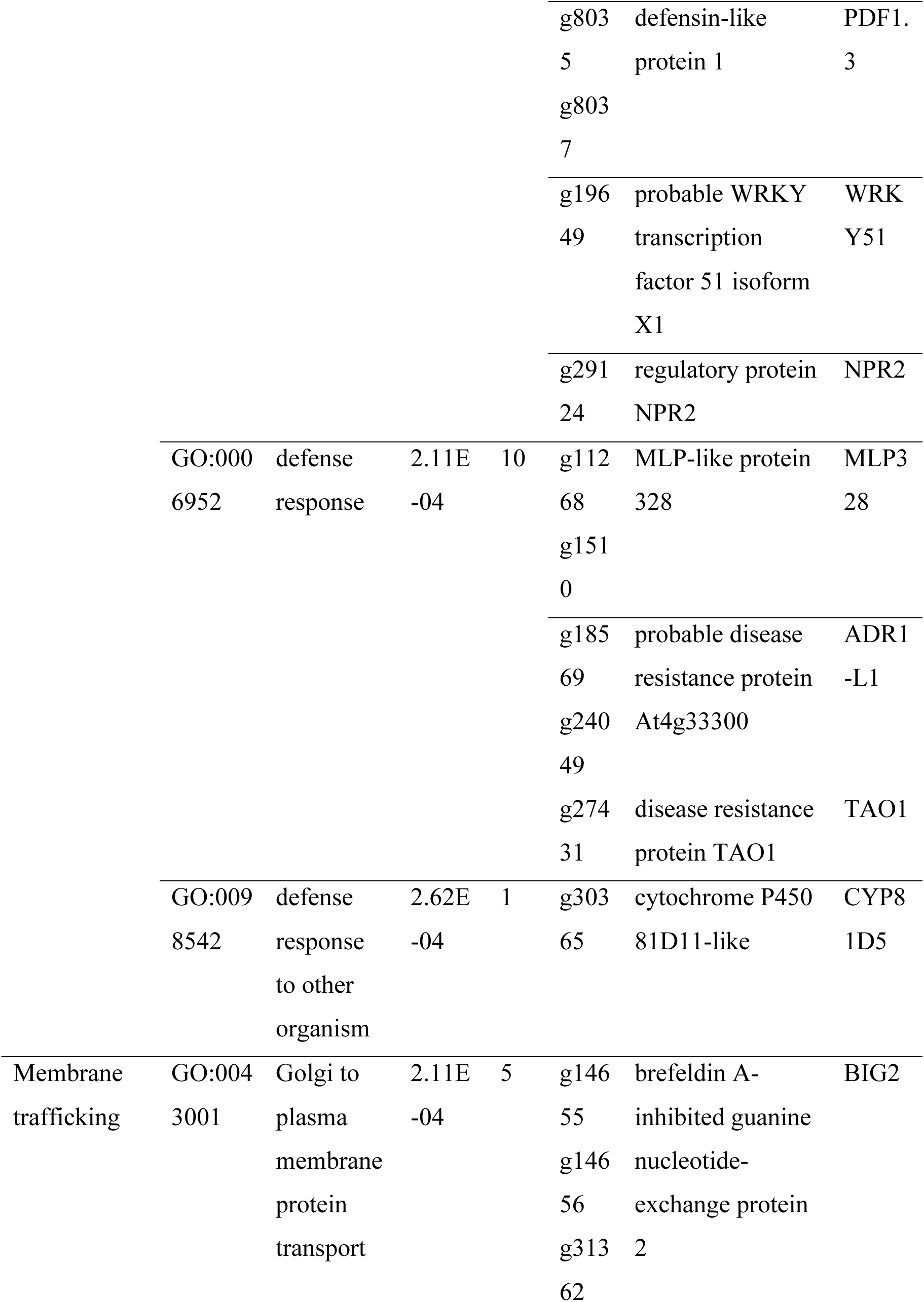
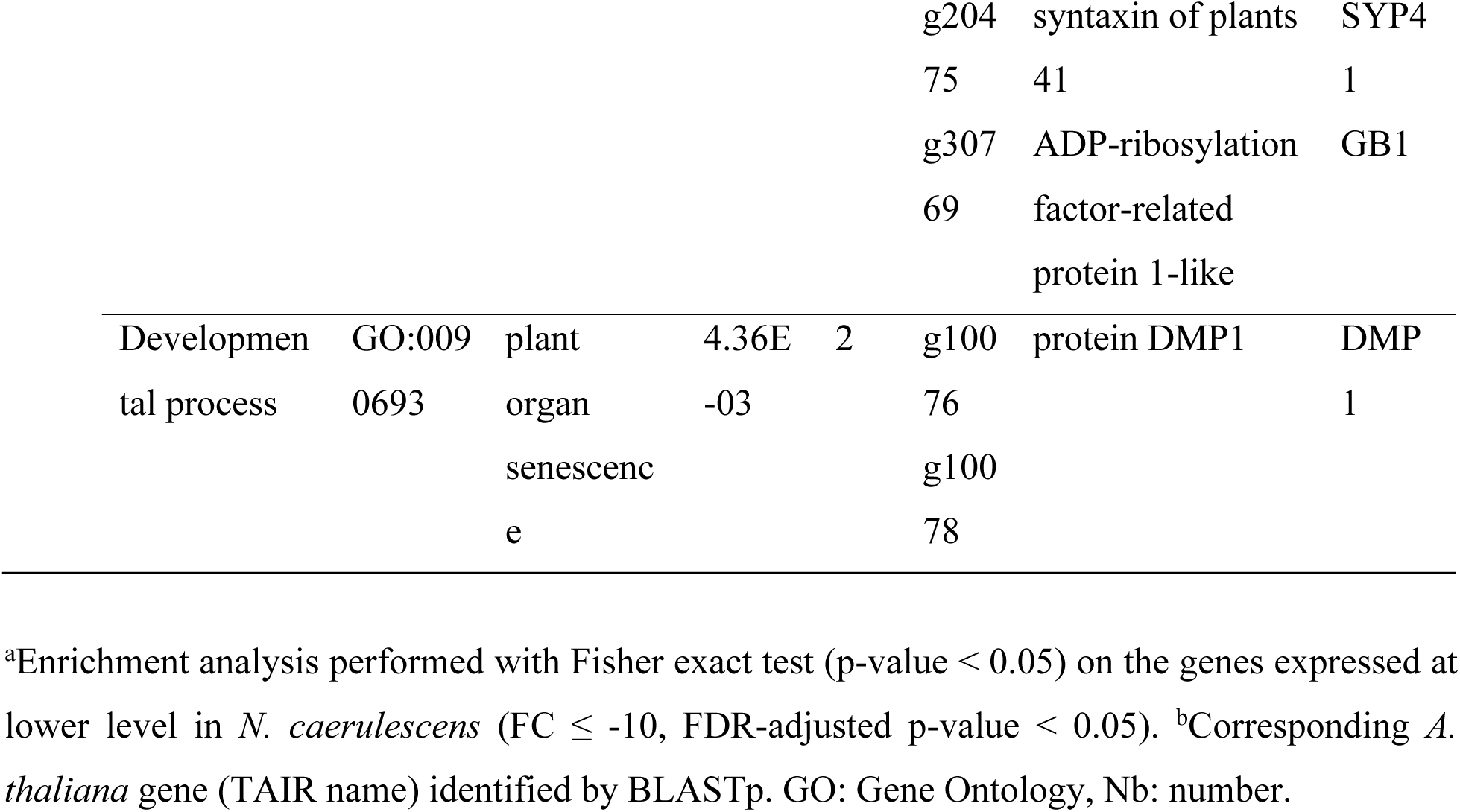
Enrichment analysis of genes exhibiting lower expression in *Noccaea caerulescens* compared to *Microthlaspi perfoliatum*^a^.

### Origin of the intraspecific variation for nickel hyperaccumulation in Noccaea caerulescens

Although high nickel accumulation capacity is observed in almost all *N. caerulescens* accessions, some accessions originating from calamine soils lack this ability (Seregin et al., 2022; Kozhevnikova et al., 2020). To determine whether the roots or the shoots are responsible for this intraspecific difference, we performed reciprocal grafting experiments between the nickel hyperaccumulating accession Cira (*Ncci*) and the non-accumulating accession La Calamine (*Nclc*) (Fig. 4, A and B). When *Nclc* shoots were grafted onto *Ncci* rootstocks, nickel accumulation in leaves (16930 ± 240 μg.g^-1^) was similar to that of self-grafted *Ncci* plants (16950 ± 510 μg.g^-1^). We did not observe any visible symptoms of nickel toxicity in the shoots (*Nclc*) of these grafted plants. In contrast, when *Ncci* shoots were grafted onto *Nclc* rootstocks, the leaves accumulated low concentrations of nickel (640 ± 40 μg.g^-1^), similar to self-grafted *Nclc* plants (580 ± 4 μg.g^-1^). These results indicate that *Nclc* roots lack at least one biological function essential for nickel hyperaccumulation in shoots, a conclusion further supported by grafting experiments between *Nclc* and *Ncfi* accessions (Supplementary Fig. S2). In these grafting experiments, we also observed a lower accumulation of nickel and zinc in the roots of *Nclc* (330 ± 40 μg nickel.g^-1^) compared to *Ncci* (4870 ± 500 μg nickel.g^-1^). However, both accessions displayed the same level of zinc accumulation in leaves (Fig. 4B).

**Figure 4.**
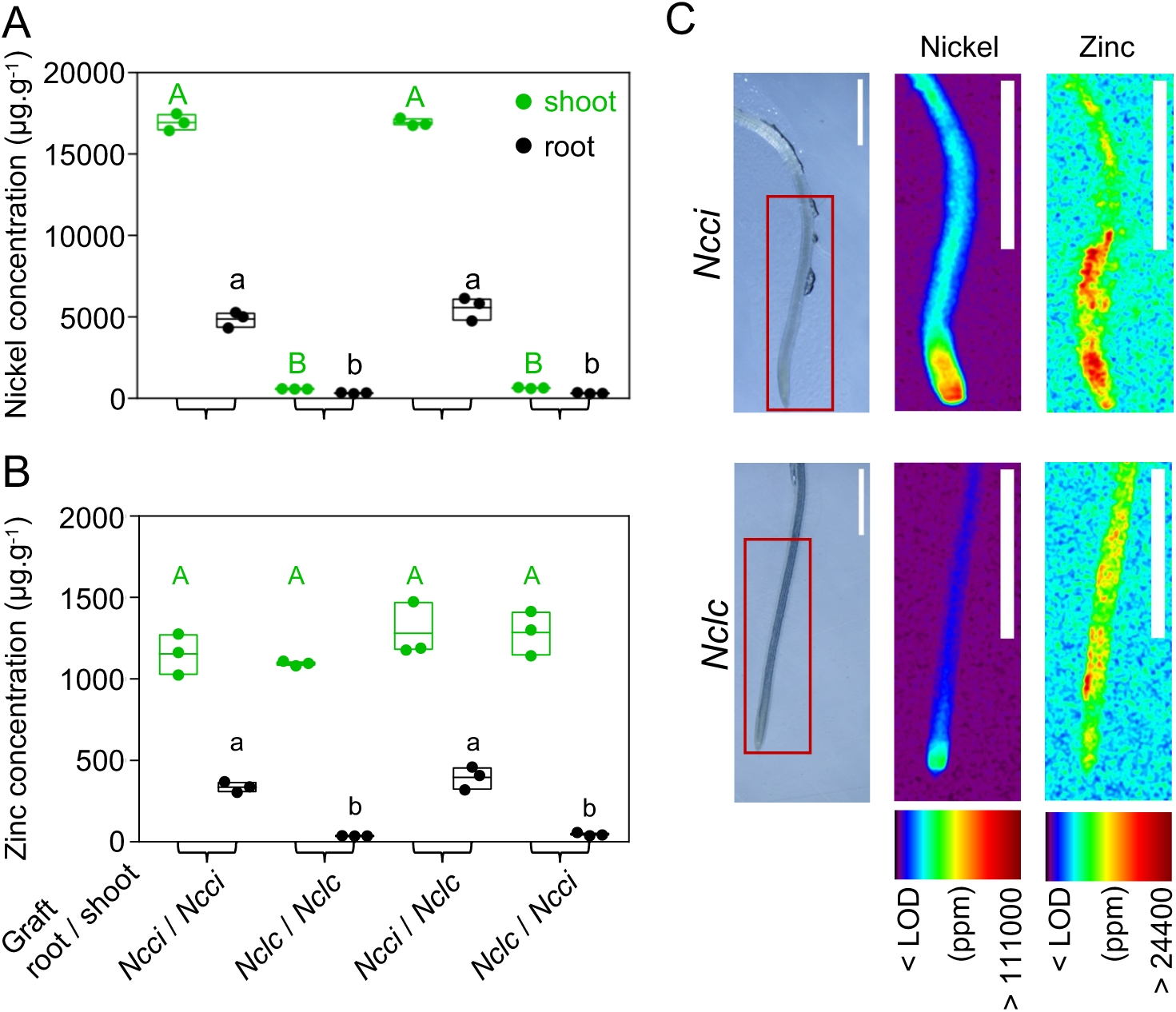
Origin of the difference in metal accumulation between *N. caerulescens* accessions. Nickel and B) zinc accumulation in shoots (green) and roots (black) of plants derived from self and reciprocal grafting of rootstock and shoot explants of *N. caerulescens* Cira (*Ncci*) and La Calamine (*Nclc*) accessions. Individual and mean metal concentration values are presented. Significant differences between the means are indicated by different letters for shoots and roots (*n* = 3 biological replicates, one-way ANOVA with Tukey’s multiple comparison test after logarithmic transformation of the data, p-value < 0.01). C) µXRF analysis of *Ncci* and *Nclc* roots for nickel and zinc semiquantitative distribution. The red square on the visible images of roots (left) represents the area analyzed by µXRF for quantification of nickel (center) and zinc (right) distribution. Metal concentrations are given in ppm. Scale bar : 1 mm. The images are representative of a set of 3 and 2 biological replicates obtained for *Ncci* and *Nclc* respectively (Supplementary Fig. S3).

To further pinpoint the defect in nickel transport in the roots of *Nclc*, we performed μXRF elemental imaging to investigate the pattern of metal accumulation in the roots of these accessions (Fig. 4C; Supplementary Fig. S3). µXRF imaging revealed a strong accumulation of nickel in the root tip and in the central part of the root of *Ncci*. In *Nclc*, nickel accumulated at a much lower level but with a similar pattern of distribution, suggesting that *Nclc* roots are affected in nickel uptake and possibly in its radial transport to the xylem. Zinc displayed a broader distribution pattern along the root axis, with a stronger accumulation in *Ncci* compared to *Nclc*. Bulk elemental analysis and µXRF imaging indicated that while zinc uptake is also affected in *Nclc* roots, it is not a limiting factor for zinc accumulation in the shoots, in our assay condition. Taken together, our data suggest that metal uptake is affected in *Nclc* roots and that this defect accounts for the inability of this accession to hyperaccumulate nickel. This result raises the hypothesis that the activity of a metal transporter, present in nickel hyperaccumulating accessions, is strongly decreased or lost in the *Nclc* accession.

### Identification of biological functions affected in N. caerulescens La calamine roots

To identify the genes underlying the missing nickel transport function in roots of La Calamine (*Nclc*), we first compared gene expression in roots of *Nclc* and of the three nickel hyperaccumulating accessions of *N. caerulescens* (i.e. *Ncfi*, *Ncbe*, *Ncpu*) to identify DE genes with a high fold change of expression (FC ≥ 10 or FC ≤ -10; Fig. 5, A and B; Supplementary Fig. S4; Supplementary Data Set S3). In each of the three comparisons, we identified between 2% and 5% DE genes (|log2FC| ≥ 3.32, FDR-adjusted p-value < 0.05, n = 3-4), but only 123 DE genes were common to all three comparisons (Fig. 5B). However, the functional annotation of the 52 genes showing strongly reduced expression in *Nclc* roots did not reveal an obvious candidate in metal transport (Supplementary Table S1). Therefore, we screened the coding sequence of genes expressed in *Nclc* roots to identify mutations that could lead to a loss-of-function of the corresponding protein. Using our RNA-Seq data and the N. caerulescens Cira (*Ncci*) genome as reference, we identified 27 977 single nucleotide polymorphism (SNP) and insertion-deletion (Indel) mutations in 9 293 genes expressed in roots of *Nclc*. After filtering out mutations also identified in *Ncfi* (28 417 SNP/Indel), *Ncbe* (26 091 SNP/Indel) and *Ncpu* (11 417 SNP/Indel), we retained 9519 SNP/Indel mutations, present in 4647 genes, unique to *Nclc* (Supplementary Data Set S4). To identify potential loss-of-function mutations, we focused on the 335 SNP/Indel mutations, affecting 221 genes, that were predicted to introduce a frameshift and/or a premature stop codon into coding sequences. Among these genes, nine genes are annotated as playing a role in nutrient and solute transport (Table 3). Interestingly, two of these genes correspond to the orthologs of the *A. thaliana* ZIP metal transporters ZIP10 (g22398) and IRT1 (g27500), which are encoded by unique genes in the *Ncci* genome. DNA resequencing of *Nclc* confirmed a 10-bp deletion in the *NcIRT1* coding sequence and a 5-bp insertion in *NcZIP10* (Fig. 5C, Supplementary Fig. S5). Both mutations are predicted to introduce a frameshift in the last exon of the corresponding genes, introducing a premature stop codon that prevent the correct translation of the last two transmembrane domains of these ZIP transporters. In addition, the resequencing the genomic DNA of the non-nickel accumulating accessions Prayon (*Ncpr*) and Plombières (*Ncpb*) revealed a mapping breakpoint at the same position in the first exon of *NcIRT1* around which paired-end reads only map as broken reads to the reference *Ncci* genome (Fig. 5C). This mapping breakpoint indicates a large DNA insertion in *NcIRT1*. Similar analyses did not reveal mutation in *NcZIP10* in these two accessions.

**Figure 5.**
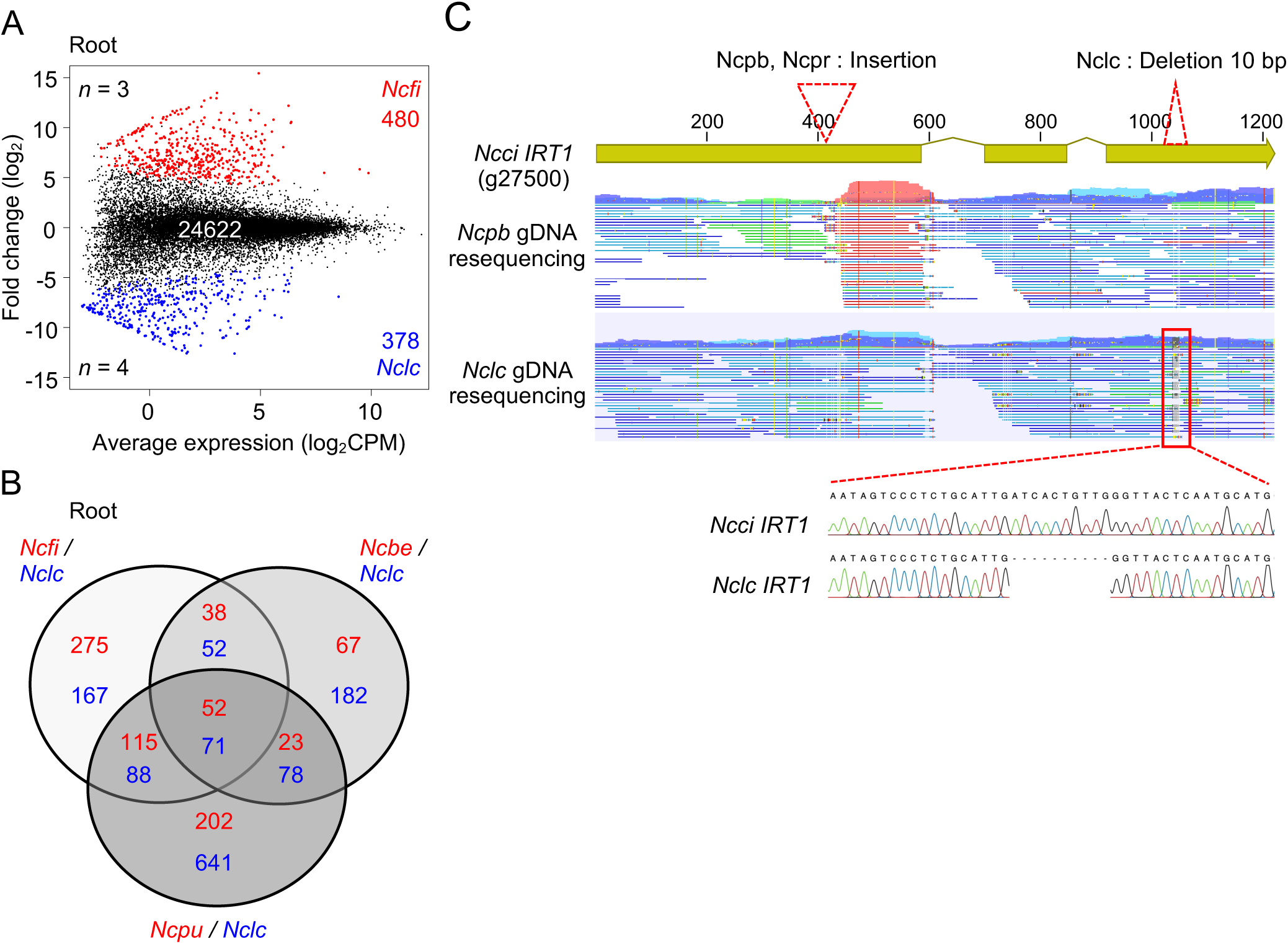
Identification of candidate genes involved in nickel transport in roots of *N. caerulescens.* A) MA plot representing the intraspecific comparative transcriptomics analysis of gene expressed in roots of *N. caerulescens* Firmiensis (*Ncfi*) and La Calamine (*Nclc*). The number of DE genes (|log_2_FC| ≥ 3.32, FDR-adjusted p-value < 0.05, *n* = 3-4 biological replicates), more expressed in *Ncfi* (red dots) or in *Nclc* (blue dots) are indicated. The comparisons of *N. caerulescens* accessions Bergenbach (*Ncbe*) and Puente basadre (*Ncpu*) with *Nclc* are presented in Supplementary Fig. S4. B) Venn diagram displaying the number of DE genes in roots between the nickel hyperaccumulating accessions (red) and *Nclc* (blue). C) Genome resequencing (gDNA) of the non-nickel accumulating *N. caerulescens* accessions Plombières (*Ncpb*), Prayon (*Ncpr*) and La Calamine (*Nclc*). In the mapping panel, paired-end reads are in blue and broken pairs are in red and green. Read mapping on the *Ncci* genome revealed a large DNA insertion at the same position in the first exon of *NcIRT1* (g27500) in *Ncpb* and *Ncpr,* and confirmed the 10 bp deletion in the third exon of *NcIRT1* in *Nclc* (also verified by Sanger sequencing).

The identification of two independent mutations predicted to affect the integrity of NcIRT1 in three distinct accessions of *N. caerulescens* unable to accumulate nickel, strongly suggests that loss of function mutations in this metal transporter account for the loss of nickel hyperaccumulation in some *N. caerulescens* accessions.

### Characterization of the metal transport activity of NcIRT1

Previous works indicated that the *A. thaliana* IRT1 transporter (AtIRT1) is able to transport nickel *in planta* (Schaaf et al., 2006; Nishida et al., 2011). To characterize the metal transport activity of NcIRT1, we performed gene functional assays in yeast. The expression of *NcIRT1*, cloned from *Ncfi*, is able to complement the growth defect of the *fet3fet4* yeast mutant in absence of iron supplementation, indicating that NcIRT1 transports iron (Fig. 6A). This result is supported by the higher accumulation of iron in yeast cells expressing *NcIRT1*, as observed with *AtIRT1* (Fig. 6C). The expression of *NcIRT1* increases nickel sensitivity and accumulation in yeast, showing that NcIRT1 is also able to transport nickel in yeast cells (Fig. 6, B and D). The higher level of zinc accumulation in yeast cells expressing *NcIRT1* compared to the empty vector control indicates that this transporter also mediates zinc uptake (Supplementary Fig. S6). Zinc was previously shown to compete with nickel for metal accumulation in *N. caerulescens* (Assunção et al., 2001; Kozhevnikova et al., 2021). Interestingly, we observed that *NcIRT1*-mediated iron and nickel accumulation in yeast is inhibited by the addition of 50 µM zinc in the culture medium (Fig. 6, B and D), suggesting that zinc competes with iron and nickel for *NcIRT1-*mediated transport. In contrast, the expression of the variant *NclcIRT1* from La Calamine neither complements *fet3fet4* nor increases nickel sensitivity (Fig. 6, A and B). Accordingly, the expression of *NclcIRT1* does not increase iron, nickel or zinc accumulation in yeast (Fig. 6, C and D, Supplementary Fig. S6). Together, these results indicate that NcIRT1 has the functional features of the metal transporter involved in nickel uptake in *N. caerulescens* and confirm that the protein encoded by the NclcIRT1 gene is nonfunctional or unstable in this expression system.

**Figure 6.**
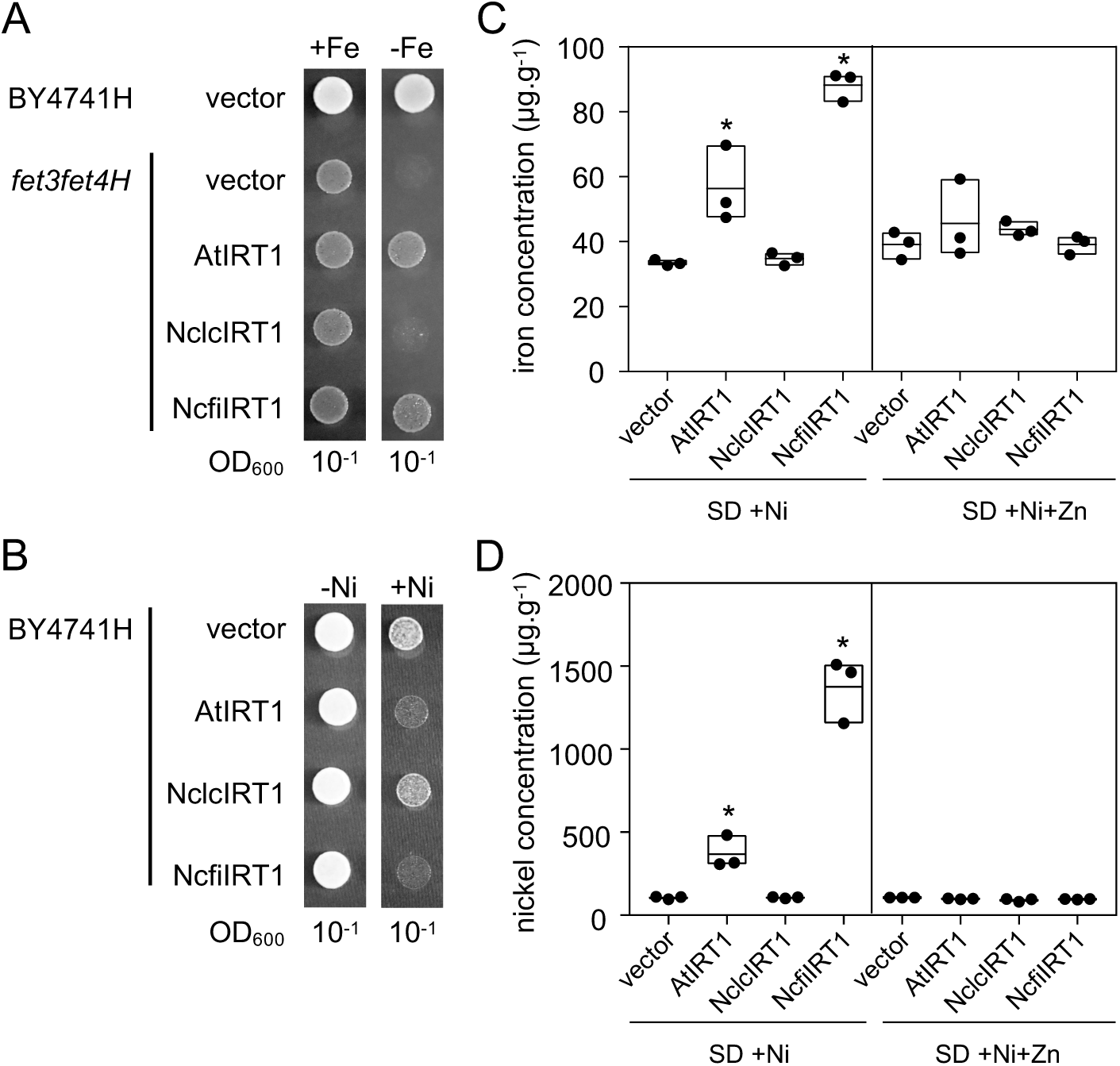
Functional characterization of NcIRT1 in yeast. A) The iron-deficient yeast mutant *fet3fet4H* was transformed with *NcIRT1* originating from the accessions Firmiensis (NcfiIRT1) and La Calamine (NclcIRT1). The BY4741H yeast strain and *fet3fet4H* transformed with the empty vector pDR195 or *A. thaliana IRT1* (AtIRT1) were used as controls. *Fet3fet4H* complementation was assessed by spotting and growing yeast transformants on SD medium (-Fe) and SD medium supplemented with 100 µM Fe-EDTA (+Fe). B) BY4741H was transformed with the same constructs and nickel tolerance was assessed by growing yeast transformants on SD medium (-Ni) or SD medium supplemented with 500 µM NiCl_2_ (+Ni). C) Iron and D) nickel accumulation were measured in BY4741H transformants grown in liquid SD medium containing 100 µM NiCl_2_ (SD +Ni) and in the same medium supplemented with 50 µM ZnSO_4_ (SD +Ni+Zn). The data represent individual measurements as well as the mean, the minimum, and the maximum values. Asterisks indicate significant differences compared to the control (vector) mean (Welch’s t-test after logarithmic transformation of the data, p-value < 0.01, n=3).

### NcIRT1 is constitutively expressed in the epidermis of N. caerulescens roots

Our transcriptomic analysis indicated that *NcIRT1* (g27500) is expressed in roots of *N. caerulescens* in the presence of iron and is not induced by the addition of nickel in the nutrient solution (Supplementary Data Set S2 and S3). To further characterize the expression pattern of *NcIRT1*, we expressed the *GUS* reporter gene under the control of the 3.8 kb genomic region located upstream of *NcIRT1*, in *N. caerulescens* hairy roots (Fig. 7). The analysis of 15 independent transformed lines showed that the *GUS* gene is expressed in the differentiation and mature zones of roots in the presence of iron in the culture medium. We did not observe the activity of the *NcIRT1* promoter in the meristematic and elongation zones of roots (Fig. 7, A and B). At higher magnification and using root transversal sections, we observed that the *NcIRT1* promoter drives the expression of *GUS* in the epidermal trichoblast cells (Fig. 7, D and E). These results show that *NcIRT1* is expressed in the epidermis of *N. caerulescens* root cells independently of the presence of iron. Together with its ability to transport nickel, and its loss of function in *N. caerulescens* accessions that do not accumulate nickel, the constitutive expression of *NcIRT1* in root epidermal cells strongly support the role of NcIRT1 in nickel uptake in hyperaccumulating accessions of *N. caerulescens*.

**Figure 7.**
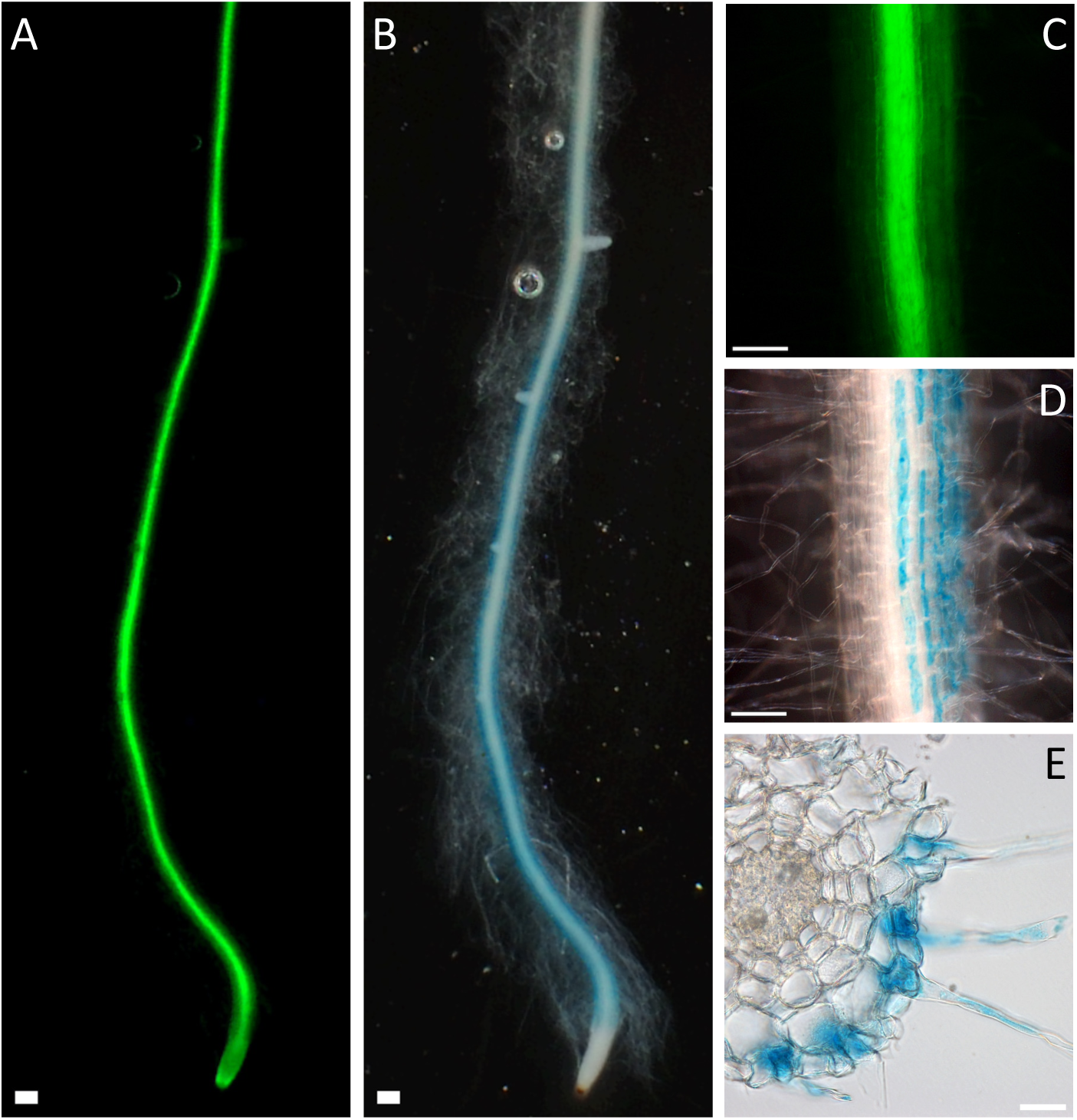
Pattern of *NcIRT1* expression in *Noccaea caerulescens* hairy roots. The *GUS* reporter gene (blue color) was expressed under the control of the *NcIRT1* promoter in *N. caerulescens* Cira transgenic hairy root lines. In these lines, the green fluorescent protein GFP, expressed under the control of the 35S promoter, was used as a transformation marker. A and B) Transgenic root imaged with A) fluorescence microscopy and B) bright field microscopy at low magnification (scale bar=100 µm). C and D) Transgenic root imaged at higher magnification (scale bar=50 µm). E) Transversal section of a transgenic root imaged with bright field microscopy (scale bar=20 µm).

### The expression of NcIRT1 partially restore nickel accumulation in N. caerulescens La Calamine

We then tested whether the expression of a functional version NcIRT1 is sufficient to restore nickel hyperaccumulation in the *N. caerulescens* non-accumulating accession La Calamine (*Nclc*). To this aim, we expressed a tagged version of *NcIRT1* from the Firmiensis accession (NcIRT1- TagRFP) under the control of the 35S promoter in *Nclc* roots. We first confirmed that the fusion protein NcIRT1-TagRFP is functional by complementing the *A. thaliana irt1* mutant (Supplementary Fig. S7). Using *Rhizobium rhizogenes*-mediated transformation, we generated 45 independent composite *Nclc* transgenic lines, 29 of which expressed NcIRT1-TagRFP at various levels as deduced from TagRFP fluorescence intensity (*Nclc*[TagRFP]^+^). The remaining 16 lines did not visibly express NcIRT1-TagRFP (*Nclc*[TagRFP]^-^) and were used as negative control. Confocal microscopy imaging of *Nclc*[TagRFP]^+^ lines showed that NcIRT1-TagRFP localized to the plasma membrane irrespective of the presence of nickel in the culture medium (Fig. 8A). Elemental analysis revealed that *Nclc*[TagRFP]^+^ lines accumulate approximately two-fold more nickel in shoots (1060 ± 330 μg.g^-1^) than *Nclc*[TagRFP]^-^ lines (480 ± 60 μg.g^-1^; Fig. 8B). We further showed that nickel accumulation in shoots of *Nclc*[TagRFP]^+^ lines is positively correlated (r^2^ = 0.87, p-value < 0.01) with the expression level of *NcIRT1-TagRFP* in roots as quantified by qRT-PCR (Fig. 8C). However, the accumulation of nickel in shoots of *Nclc*[TagRFP]^+^ lines does not reach the level measured in shoots of wild-type *N. caerulescens* Cira (*Ncci*; 3560 ± 600 μg.g^-1^) or *Ncci* hairy root control lines (4330 ± 1862 μg.g^-1^). The expression of *NcIRT1-TagRFP* also increases nickel accumulation in roots of transformed *Nclc* lines, although this accumulation is not directly correlated with the level of *NcIRT1-TagRFP* expression (Fig. 8, D and E). These results show that the expression of a functional NcIRT1 transporter in *Nclc* roots increases nickel uptake and indicate that this step is a limiting factor for nickel hyperaccumulation in shoots. However, the ectopic expression of NcIRT1 in *Nclc* rootstock is not sufficient to restore the level of nickel accumulation observed in shoots of nickel hyperaccumulating accessions.

**Figure 8.**
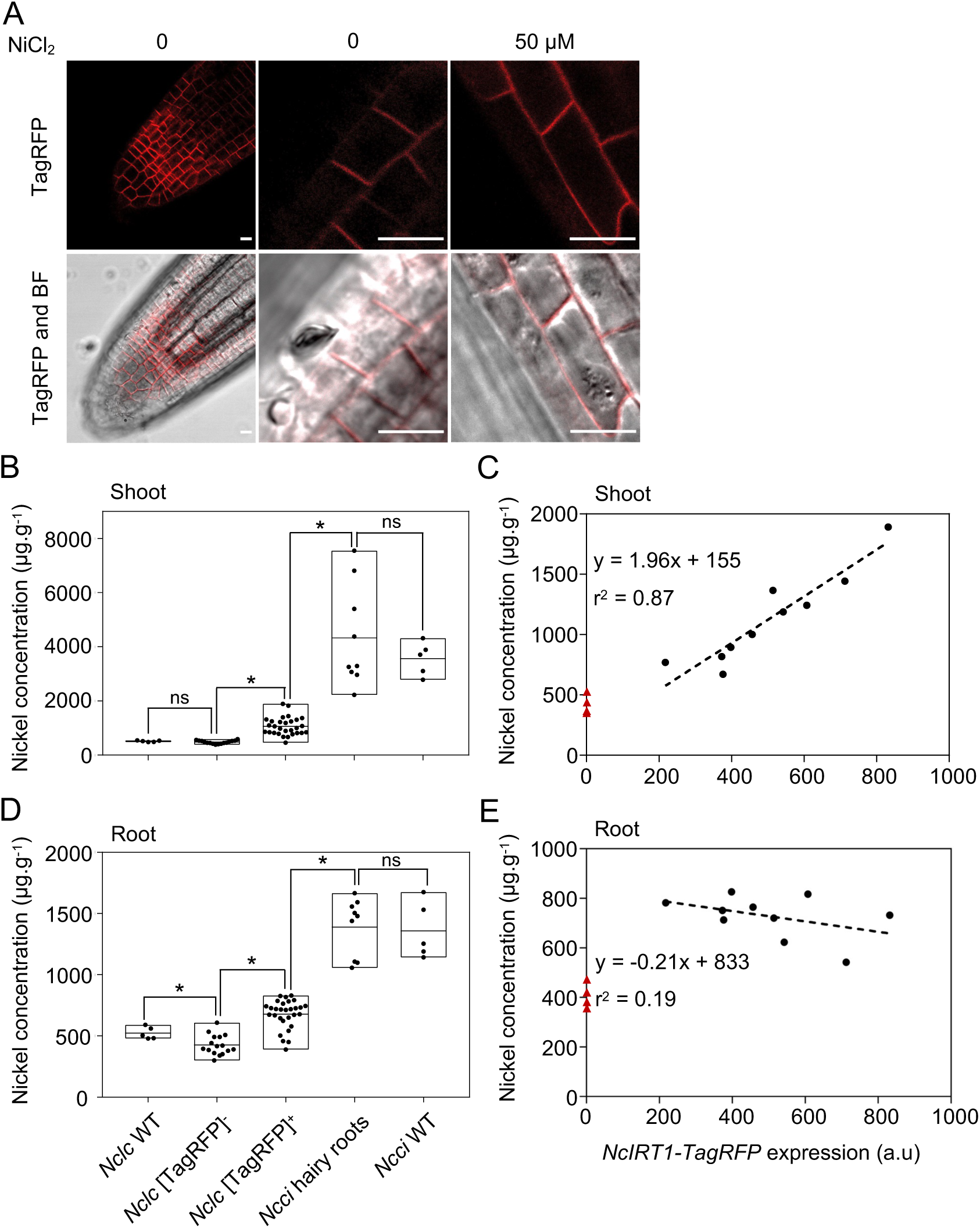
Expression of NcIRT1-TagRFP in *N.caerulescens* La Calamine (*Nclc*). A) *Nclc* hairy root lines expressing NcIRT1-TagRFP were grown *in vitro* without and with 50 µM NiCl_2_. Root were imaged at the root tip by confocal microscopy to visualize the TagRFP signal (red channel). BF: Bright-field. Scale bar, 10 µm. Images are representative of *n* ≥ 3 biological replicates. B) Nickel accumulation in shoots of *Nclc* and Cira (*Ncci*) accessions (*n* = 5 each), *Ncci* hairy roots lines transformed with empty vector (*n* = 9), and *Nclc* lines expressing NcIRT1-TagRFP (*Nclc*[TagRFP]^+^, *n* = 29) or not (*Nclc*[TagRFP]^-^, *n* = 16). Plants were grown in hydroponic culture containing 50 µM NiCl_2_ for 4 weeks. The data represent individual measurements and corresponding box (max., min., average). Asterisks indicate significant differences (Welch’s t-test after logarithmic transformation of the data, p-value < 0.01, n=3). C) Correlation analysis (Pearson test) of nickel accumulation in shoots and *NcfiIRT1-TagRFP* expression quantified by qRT-PCR in *Nclc*[TagRFP]^+^ lines (*n* = 10, black dots) and (*Nclc*[TagRFP]^-^ lines (n=4, red triangles). *NcfiIRT1-TagRFP* expression was normalized with *Nc6PGDH*. Linear regression on *Nclc*[TagRFP]+ data and r squared are indicated. D and E) same as B) and C) but for nickel accumulation in roots.

## Discussion

*Noccaea caerulescens* has been proposed for several years as a genetic model to study metal hyperaccumulation. However, molecular and genetic studies on this species have been limited due to the lack of a high-quality genome assembly. In this work, we obtained a chromosome-level assembly of the *N. caerulescens* genome and used this new genome assembly as a reference to identify the molecular mechanisms associated with the evolution of nickel hyperaccumulation in *N. caerulescens*. Our inter and intraspecific transcriptomic comparisons suggest that nickel hyperaccumulation evolved from the high and constitutive expression of only few genes involved in the transport of metals including NcIREG2. In addition, we provide genetic and functional evidence that the metal transporter NcIRT1 is involved in nickel uptake and accumulation, and that loss-of-function mutations in this gene are responsible for the loss of the nickel hyperaccumulation trait in some *N. caerulescens* accessions.

### A high-quality genome assembly for Noccaea caerulescens

Using long-read sequencing, we assembled the genome of the *N. caerulescens* accession Cira into seven large scaffolds corresponding to the chromosomes of this species (Figure 1, Table 1). This assembly of 254 Mb is close to the total genome size of 304 Mb calculated by k-mer analysis, and the estimated genome size of 267 Mb obtained by flow cytometry (Mandáková et al., 2015). The analysis of this assembly revealed that the *N. caerulescens* genome contains approximately 60% of repetitive and transposable elements (RE/TE). This high proportion of RE/TE is significantly higher than in *A. thaliana* (∼20%) and similar to the proportion of RE/TE found in *Thlaspi arvense* (Contreras-Garrido et al., 2024).The presence of long stretches of repetitive sequences likely affected the previous assembly of the *N. caerulescens* genome using short-read sequencing (Kiefer et al., 2019). The overall structure of our assembly is compatible with the structure of the *N. caerulescens* genome inferred from cytogenetic analysis (Mandáková et al., 2015). However, we observed few differences between these two proposed structures. For example, our assembly suggests that the chromosomes 3 and 5 are acrocentric (Figure 1), which was not observed using cytogenetic analysis. We also observed differences in the proposed gene arrangement in the lower arm of chromosome 5 and around the *IRT1* locus on chromosome 7. The genome of the Cira accession, which was not analyzed by cytogenetics, may contain specific structural variations that are not present in other accessions. In addition, chromosome painting may not have captured small structural variations present in the *N. caerulescens* genome. On the other hand, we cannot exclude that the presence of long stretches of RE/TE affected the scaffolding of the *N. caerulescens* contigs using the chromatin capture strategy. The use of ultra-long read sequencing, or optical map would help to further improve the quality of the *N. caerulescens* genome assembly (Garg et al., 2024). Even if it may be further improved, the assembly obtained in this work represents a significant improvement of the *N. caerulescens* genome sequence and structure and provides a valuable tool for future genomic and transcriptomic studies in this model species.

### Genetic pathways associated with nickel hyperaccumulation

Our RNA-Seq analysis revealed that the *N. caerulescens* ultramafic-adapted accession Firmiensis is essentially insensitive at the transcriptomic level to the presence and accumulation of nickel in roots and shoots (Fig. 2, A and B). In contrast, in the related non-accumulator species *Microthlaspi perfoliatum,* subtoxic concentration of nickel induces a transcriptomic response (Fig. 2, A and B; Supplementary Data Set 2), reminiscent of the iron deficiency response as previously observed in *A. thaliana* (Lešková et al., 2020). This result suggests that nickel hyperaccumulation in *N. caerulescens* is a constitutive trait rather than an adaptive trait induced in response to nickel. We thus performed interspecific transcriptomic comparisons between *N. caerulescens* and *M. perfoliatum* to identify genes whose expression is associated with nickel hyperaccumulation. This analysis revealed that several genes involved in defense responses against pathogens are less expressed in the shoot of *N. caerulescens* (Table 2). This reduction of the expression of defense genes likely illustrates a trade-off between induced disease resistance and elemental defense in *N. caerulescens*, thus supporting the elemental defense theory proposed as a selective advantage conferred by nickel hyperaccumulation (Hörger et al., 2013). This analysis also reveal that several genes involved in membrane trafficking are expressed at a lower level in *N. caerulescens*. Studies in yeast have previously showed that loss of function mutation in several genes involved in membrane trafficking confer resistance to nickel (Arita et al., 2009; Ruotolo et al., 2008). The regulation membrane trafficking is recognized as an important mechanism for the control of plant metal transporter activity (Ivanov and Vert, 2021). Therefore, the reduction of the expression of genes involved in membrane trafficking in *N. caerulescens* could be directly linked to the unusual regulation of metal homeostasis in metal hyperaccumulators.

In contrast, our analysis did not reveal any enrichment of differentially expressed (DE) genes involved in the regulation of metal homeostasis. This result was surprising because previous transcriptome analyses have suggested that zinc and nickel hyperaccumulation is associated with the high expression of a set of more than 30 genes involved in metal homeostasis (Meier et al., 2018; Hanikenne and Nouet, 2011; Halimaa et al., 2014a). The results of these studies are difficult to compare because they use different species or accessions and use different fold-change and statistical criteria to identify DE genes. In this study, we also used three ultramafic-adapted accessions of *N. caerulescens* belonging to different genetic units to represent the nickel hyperaccumulator group (Gonneau et al., 2017), thus excluding accession-specific DE genes that could be representative of local adaptation. Our analysis identified only three genes encoding for orthologs of metal transporters, namely *NcHMA4*, *NcHMA3* and *NcIREG2*, that are reliably expressed at higher level in nickel hyperaccumulating accessions of *N. caerulescens* compared to *M. perfoliatum*. These results suggest that nickel hyperaccumulation has evolved from the high and constitutive expression of a limited set of metal transporters.

### Metal transporters involved in nickel hyperaccumulation

The gene encoding the IREG/FPN transporter NcIREG2 (g26066) is significantly more highly expressed in leaves of *N. caerulescens* compared to *M. perfoliatum* (Supplementary Data Set S3). This result confirms previous studies showing that the high expression of IREG/FPN genes is repeatedly associated with nickel hyperaccumulation in a wide diversity of species (Meier et al., 2018; García de la Torre et al., 2021; Halimaa et al., 2014a; Merlot et al., 2014). Both NcIREG2 and its ortholog in *A. thaliana*, AtIREG2, are able to transport nickel and localize to the vacuolar membrane (Schaaf et al., 2006; García de la Torre et al., 2021). Furthermore, the silencing of *NcIREG2* in the roots of transgenic *N. caerulescens* plants correlates with a reduction in nickel accumulation (García de la Torre et al., 2021). Taken together, these results strongly suggest that the high expression of *NcIREG2* in leaves is essential for the storage of nickel in the vacuole of epidermal cells, as observed by elemental imaging in *Noccaea* species (Küpper et al., 2001; van der Ent et al., 2019; do Nascimento et al., 2021).

Our data also show that the P1B-type ATPase genes *NcHMA3* and *NcHMA4* are highly expressed in *N. caerulescens* compared to *M. perfoliatum*. The high expression of *NcHMA4* and *NcHMA3* has previously been associated with the ability of *N. caerulescens* to hyperaccumulate zinc (Bernard et al., 2004; Papoyan and Kochian, 2004; Ueno et al., 2011; van de Mortel et al., 2006; Ó Lochlainn et al., 2011; Halimaa et al., 2014a), a trait found in all *N. caerulescens* accessions studied so far (Kozhevnikova et al., 2020). In particular, the high expression of the ortholog of NcHMA4 in *A. halleri*, AhHMA4, has been shown to be necessary and sufficient for efficient root-to-shoot translocation of zinc and has been crucial for the evolution of zinc tolerance and hyperaccumulation in this species (Hanikenne et al., 2008). Therefore, it is surprising to observe the conservation of high *NcHMA4* expression in three *N. caerulescens* accessions originating from ultramafic soils that are relatively poor in zinc. This result suggests that high *NcHMA4* expression may confer a selective advantage for the growth of *N. caerulescens* in these particular edaphic conditions. One hypothesis is that the high expression of *NcHMA4* favors zinc distribution in the shoots of *N. caerulescens* in a context of high nickel concentration competing with zinc. A second hypothesis is that high *NcHMA4* expression is directly involved in nickel tolerance and accumulation in *N. caerulescens*. Further experiments will be required to measure the activity of NcHMA4 towards nickel (Eren and Argüello, 2004; Mishra et al., 2017). Even if the high expression of *NcHMA4* participate in the hyperaccumulation of nickel in *N. caerulescens*, this is unlikely to represents a highly conserved or convergent mechanism, as high expression of HMA4 orthologs has not been observed in other nickel hyperaccumulating species (Meier et al., 2018; García de la Torre et al., 2021). Other metal transporters of the IREG/FPN family localizing to the plasma membrane have been proposed to mediate nickel loading in the xylem and root-to-shoot translocation (Morrissey et al., 2009; González et al., 2024). However, the corresponding gene in *N. caerulescens, NcIREG1* (g15128), is not highly expressed compared to *M. perfoliatum* (Supplementary Data Set S3).

### Role of NcIRT1 in nickel hyperaccumulation

Interspecific transcriptome comparison between *N. caerulescens* and *M. perfoliatum* did not reveal candidate metal transporters involved in nickel uptake in roots. However, intraspecific reciprocal grafting between rootstocks and shoots of Cira and La Calamine accessions clearly indicated that nickel hyperaccumulation requires at least one important function in the roots of *N. caerulescens* that is missing in La Calamine (Fig. 4A; Supplementary Fig. S2). Intraspecific transcriptome comparison did not reveal any candidate gene involved in metal transport that would be expressed at much lower levels in La Calamine roots compared to the nickel hyperaccumulating accessions (Fig. 5, A and B; Supplementary Table S3; Supplementary Fig. S4). In contrast, we identified frameshift mutations in the two related ZIP family transporters *NcIRT1* (g27500) and *NcZIP10* (g22398) in La Calamine that, at least in the case of NcIRT1, abolish their metal transport activity (Fig. 5C; Fig. 6; Supplemental Fig. S5). The 10-base pair deletion identified in the last exon of La Calamine *NcIRT1* is different from the previously described frameshift mutation in the first exon of this gene (Halimaa et al. 2019), suggesting that multiple *NcIRT1-*deficient alleles coexist in this population. In addition, we identified a large DNA insertion in *NcIRT1* in the Belgian accessions Prayon and Plombières (Fig. 5C), which are geographically and genetically close to La Calamine, and do not accumulate nickel (Gonneau et al., 2017; Kozhevnikova et al., 2020; Seregin et al., 2022). This DNA insertion is compatible with the truncated *NcIRT1* transcript previously observed in the Prayon accession (Plaza et al., 2007). These results indicate that there are several independent loss-of-function mutations in *NcIRT1* associated with a reduction in nickel accumulation, strongly supporting the role of *NcIRT1* in nickel hyperaccumulation. In contrast, loss-of-function mutation in *NcZIP10* does not seem to be recurrently associated with the loss of nickel hyperaccumulation in *N. caerulescens*.

Heterologous expression in yeast confirmed that NcIRT1 is able to transport nickel (Fig. 6, B and D), like its ortholog in *A. thaliana* AtIRT1 (Nishida et al., 2011). Interestingly, the addition of zinc abolishes NcIRT1-mediated nickel transport in yeast, suggesting that NcIRT1 has a higher affinity for zinc than for nickel (Fig. 6D). Physiological analyses have previously shown that zinc has a strong inhibitory effect on nickel uptake and accumulation in *N. caerulescens, N. tymphea* and *N. alpestris,* but that this inhibitory effect is less pronounced in the La Calamine accession of *N. caerulescens* (Taylor and Macnair, 2006; Deng et al., 2014; Assunção et al., 2001, 2008). Therefore, the apparent specificity of NcIRT1 for nickel and zinc is consistent with the properties of the nickel uptake system of several *Noccaea* species hyperaccumulating nickel.

Both RNA-Seq data and promoter analyses showed that *NcIRT1* is expressed in roots of *N. caerulescens* even in the presence of iron in the culture medium (Fig. 7; Supplementary Data Set S2). This result contrast with the metal-sensitive species *A. thaliana*, in which the expression of the orthologous *AtIRT1* gene is only detected in roots under iron-deficient conditions (Vert et al., 2002). High expression of both *NcIRT1* and *NcZIP10*, was associated with nickel accumulation when the transcriptome of the ultramafic accession Monte Prinzera was compared with those of the calamine accessions Ganges and La Calamine (Halimaa et al., 2014a). High expression of the ortholog of IRT1 was also associated with nickel hyperaccumulation in the Asteraceae species *Senecio coronatus* (Meier et al., 2018). These results suggest that the constitutive and high expression of IRT1 orthologs is important to support the efficient uptake of nickel, and likely represent a convergent step in the evolution of nickel hyperaccumulation in different plant species. However, our interspecific transcriptome comparison did not reveal a higher expression of *NcIRT1* in *N. caerulescens* compared to *M. perfoliatum* (Supplementary Data Set S3), suggesting that the high and constitutive expression of *NcIRT1* may have evolved early in the Coluteocarpeae tribe, before the divergence of the *Noccaea* and *Microthlaspi* genera.

To provide direct genetic evidence for the role of *NcIRT1* in nickel hyperaccumulation in *N. caerulescens,* we first attempted to knock-down its expression in a nickel-hyperaccumulating accession using amiRNA (García de la Torre et al., 2021). However, we were unable to generate *N. caerulescens* hairy root lines in which *NcIRT1* expression was efficiently reduced. We thus ectopically expressed an active tagged form of NcIRT1 (NcIRT1-TagRFP) in the roots of the La Calamine accession. In these composite transgenic plants, NcIRT1 localizes at the plasma membrane of root cells and high concentrations of nickel do not affect this localization (Fig. 8A). Ectopic expression of *NcIRT1* in La Calamine roots results in increased accumulation of nickel in shoots. This result indicates that the activity of NcIRT1 is a limiting factor for nickel accumulation in the La Calamine accession (Fig. 8, B and C). However, the ectopic expression of *NcIRT1* in La Calamine is not sufficient to restore nickel accumulation to similar level as observed in the nickel hyperaccumulating accession Cira (Fig. 8B). It is possible that the strong and ectopic expression of *NcIRT1* using the 35S promoter affects the radial transport of nickel from epidermal cells to the root vasculature and subsequent translocation to the leaves (Barberon et al., 2014; Dubeaux et al., 2015). Moreover, a functional NcZIP10 may be also required to fully restore the uptake and/or the radial transport of nickel in *N. caerulescens* La Calamine. Further experiments will be required to investigate the role of *NcZIP10* in nickel hyperaccumulation.

### Evolution of nickel hyperaccumulation in N. caerulescens

The results presented here support an emerging scenario for the evolution of the nickel hyperaccumulation trait in *N. caerulescens*, involving both gain and loss of function in genes involved in metal transport. The high expression of the vacuolar IREG/FPN transporter in leaves of three nickel hyperaccumulator accessions from different genetic origin suggests that this high expression represents an essential step in the evolution of nickel hyperaccumulation in *N. caerulescens.* As in other species, the evolution of nickel hyperaccumulation in *N. caerulescens* most certainly occurred either as a result of or in parallel with the adaptation to ultramafic soils, also known as the serpentine syndrome, which involves an increased tolerance to nickel (Kazakou et al., 2008; Konečná et al., 2020). We propose that the increased expression of the vacuolar nickel transporter NcIREG2 in leaves represents a founding event in the adaptation of *N. caerulescens* to ultramafic soils, increasing both nickel tolerance and the capacity of this species to accumulate this metal (Turner et al., 2010; Konečná et al., 2021; García de la Torre et al., 2021; Arnold et al., 2016). The high and constitutive expression of NcIRT1 in the roots of *N. caerulescens* is likely to play an important role in the efficient uptake of nickel from ultramafic soils. The involvement of NcIRT1, and possibly of NcHMA4, in both nickel and zinc/cadmium hyperaccumulation may explain why these traits are conserved in nearly all *N. caerulescens* accessions, regardless of the nature of soil from which they originate. However, the pathways involved in nickel or zinc hyperaccumulation are not identical. Thus, the chronology of the evolution of these two traits in *N.* caerulescens, or more generally in the *Noccaea* genus, remains a puzzling question. If nickel hyperaccumulation appeared first and is associated to the adaptation to ultramafic soils, then we should find, in addition to *NcIREG2*, genetic traces of this adaptation in *N. caerulescens* accessions that develop on non-metalliferous or calamine soils (Turner et al., 2010; Arnold et al., 2016; Konečná et al., 2021). Future transcriptomic and genomic studies on a wider diversity of *N. caerulescens* accessions and *Noccaea* species should resolve this question.

Our finding further indicates that the loss of nickel hyperaccumulation capacity in few accessions of *N. caerulescens*, including La Calamine, Prayon and Plombières, is the consequence of the loss of NcIRT1, and potentially NcZIP10, functions. The loss of NcIRT1 activity also likely limits the zinc and cadmium uptake, which may provide a selective advantage to plants growing on calamine soils strongly enriched in these metals (Schvartzman et al., 2018; Corso et al., 2018; Halimaa et al., 2019).

Our study provides novel genomic insight into the mechanisms driving the evolution of metal hyperaccumulation in *N. caerulescens*. The high expression of *NcHMA4* in *N. caerulescens* is associated with its tandem duplication on chromosome 7 in the Cira accession (Fig. 1). Previous works have shown that the number of *NcHMA4* tandem copies varies from 2 to 4 in the genome of various accessions of *N. caerulescens* (Ó Lochlainn et al., 2011; Iqbal et al., 2013; Craciun et al., 2012). In comparison, the ortholog of *HMA4* is present as a single copy gene in the genome of *Microthlaspi erraticum*, a non-accumulator species closely related to *M. perfoliatum* (Mishra et al., 2020). Tandem triplication and cis-regulatory modifications were shown to be involved in the high expression of *AhHMA4* and the evolution of zinc hyperaccumulation in *A. halleri* (Hanikenne et al., 2008). In contrast, the high and constitutive expression of *NcIREG2* and *NcIRT1* does not seem to be the result of gene duplication, at least in the Cira accession (Fig. 1). In these cases, we propose that modifications in cis-regulatory elements of the promoters of these genes are responsible for their high activity in *N. caerulescens*. Transposable elements (TE) are known drivers of the evolution of genomes, affecting gene expression or activity through various mechanisms, such as insertion in coding sequences, gene duplication, methylation of surrounding DNA, or insertion of cis-elements in promoter regions (Quesneville, 2020; Lisch, 2013; Dubin et al., 2018). The genome of *N. caerulescens* is rich in TE (Table 1) and ancient insertion of TE in the promoter regions of *NcIRT1* and *NcIREG2* might have brought cis-regulatory elements leading to the high and constitutive expression of these genes in *N. caerulescens*. Additionally, the large DNA insertion observed in the *NcIRT1*coding region in Prayon and Plombières accessions is consistent with a recent TE insertion. Further analyses of TE insertions and activity in *N. caerulescens* is necessary to confirm their role in the evolution of metal hyperaccumulation.

## Materials and methods

### Plant material and growth condition

In this study, we used several *Noccaea caerulescens* accessions adapted to ultramafic soils and hyperaccumulating nickel: Firmiensis (*aka* Puy de Wolf, *Ncfi*; N 44°33.217 E 2°18.320, France), Bergenbach (*Ncbe*; France; provided by Dr Claude Grison), Puente Basadre (*Ncpu*; N 42°49.885, W 8°00.196, Spain), and Cira (*Ncci*; N 42°46.578, W 8°21.037; Spain); the calamine soil adapted accession La Calamine (*Nclc*; Belgium; line WUR#42), and the closely related non-accumulator species *Microthlaspi perfoliatum* (tetraploïd accession) adapted to limestone (*Mper*; N 44°23.212 E 0°49.015, France). Seeds were scarified with sandpaper and surface sterilized with 70 % ethanol, 0.05 % sodium dodecyl sulphate (SDS) for 5 min and then with 0.9 % hypochlorite (commercial bleach), 0.05 % SDS for 10 min. Seeds were then rinsed 3 times in sterile MilliQ H2O and stratified in water for 1 week at 4 °C in the dark. Seeds were germinated on half-strength Murashige and Skoog medium (1⁄2 MS), 1% agar medium, pH 5.8 and grown in a cabinet at 25 °C with 16 h light. Seedlings were then transferred to hydroponic culture and grown as previously described (**García de la Torre et al., 2021**) in a modified Hoagland’s solution containing 2 mM MES-KOH, pH 5.7, 2 µM ZnSO4, 20µM Fe-HBED and supplemented or not with NiCl2 as indicated.

### Noccaea caerulescens genome sequencing, assembly, and annotation

Fresh leaves were collected from a second self-crossed generation of *N. caerulescens* Cira plants grown in hydroponic culture for 8 weeks. Leaves were frozen in liquid nitrogen and DNA extracted with the Nucleobond High Molecular Weight DNA kit (Macherey-Nagel) according to the manufacturer’s protocol. The DNA library and sequencing were performed according to the PacBio instructions for SMRTbell Express Template Prep Kit 2.0. At each step, DNA was quantified using the Qubit dsDNA HS Assay Kit (Life Technologies). DNA purity was tested using nanodrop (Thermofisher) and size distribution assessed using the Femto pulse Genomic DNA 165 kb Kit (Agilent). Purification steps were performed using AMPure PB beads (PacBio). 15 µg of DNA was purified and sheared at 15 kb using the Megaruptor1 system (Diagenode). Using SMRTbell Express Template prep kit 2.0, a single strand overhangs removal, a DNA and END damage repair step were performed on 10 µg of sample. Then blunt hairpin adapters were ligated to the library. The library was treated with an exonuclease cocktail to digest unligated DNA fragments. A size selection step using a 12 kb cutoff was performed on the BluePippin Size Selection system (Sage Science) with “0.75 % DF Marker S1 3-10 kb Improved Recovery” protocol. Using Binding kit 2.0 and sequencing kit 2.0, the primer V2 annealed and polymerase 2.0 bounded library was sequenced by diffusion loading onto 1 SMRTcell on a Sequel2 instrument at 50 pM with a 4 hours pre-extension and a 30 hours movie. Genome characteristics were estimated with GenomeScope v. 2.0 (Vurture et al., 2017) using k-mer profile generated by Jellyfish (Marçais and Kingsford, 2011). HiFi reads were assembled with Hifiasm (version 0.15.5) at 30X coverage (Cheng et al., 2021). Assembly quality was estimated with BUSCO (Manni et al., 2021). To further scaffold genomic contigs, an Omni-C library was prepared using Dovetail Omni- C™ Kit according to the manufacturer’s protocol. Briefly, 1.5g of young leaves from *N. caerulescens* Cira were frozen in liquid nitrogen before the chromatin was crosslinked in the nucleus. Fixed chromatin was digested with DNase I and then extracted. Chromatin ends were repaired and ligated to a biotinylated bridge adapter followed by proximity ligation of adapter containing ends. After proximity ligation, crosslinks were reversed, and the DNA purified from proteins. Sequencing library was generated using Illumina-compatible adapters. Biotin-containing fragments were isolated using streptavidin beads before PCR enrichment of the library. The library was sequenced on an Illumina MiSeq platform to generate ∼5 million 2 x 150 base paired-end reads. The paired Omni-C files were aligned to the contigs using Juicer v. 1.5.7, scaffolded with 3d-dna v.529ccf4 using a previously described procedure (Istace et al., 2021) with parameter -r 0. Hi-C contact maps was manually curated with juicebox v.1.11.08 to produce a chromosome level genome assembly. The scaffolds were manually assigned to *N. caerulescens* chromosomes using Brassicaceae ABC gene blocks (Murat et al., 2015; Mandáková et al., 2015). The assembly (v2.3) was annotated using the OmicsBox suite v.1.4.12 (www.biobam.com/omicsbox). Repetitive elements were masked using RepeatMasker v.4.0.9 (Smit et al., 2015) with the custom repetitive elements sequence library generated with RepeatModeler v.2.0.1. Genes were annotated with the Eukaryotic Gene Finder tool based on AUGUSTUS v.3.4.0 (Hoff and Stanke, 2019), using *Arabidopsis thaliana* as a model to predict coding sequences and UTR, and *Ncfi* RNA-Seq read samples (SRX24619830, SRX24619832) as RNA-Seq hints (Dobin et al., 2013). The putative function of translated genes was annotated with Mercator4 v2.0 (Schwacke et al., 2019) and Blast2GO (Gotz et al., 2008) using Blastp (E-value ≤ 10^-5^) on Brassicaceae ref seq_protein database and InterProScan (Blum et al., 2021). The graphical representation of the assembly was performed with ShinyCircos (Yu et al., 2018).

### RNA extraction and sequencing

Total RNA from leaves and roots were extracted with the RNeasy Plant Mini kit (Qiagen) from three biological replicates of *Mper*, *Ncfi*, *Ncbe*, and *Ncpu,* and one replicate of *Nclc* grown in hydroponic culture (see above) for 7 weeks in Hoagland solution supplemented or not with 37.5 µM NiCl2. RNA quality control, cDNA libraries and sequencing using Illumina HiSeq2000 platform were performed as previously described (García de la Torre et al., 2021). We complemented our RNA-Seq dataset for La Calamine accession with previously published datasets (Blande et al., 2017): NcLc_R_Rep1 (SRR3743019), NcLc_R_Rep2 (SRR3743020) and NcLc_R_Rep3 (SRR3743021).

### Differential gene expression and enrichment analyses

Mapping of sequencing reads were performed using CLC Genomics Workbench v.22.0.1 (default parameters, length fraction = 0.5, similarity fraction = 0.875, paired reads counted as two) on the *N. caerulescens* Cira genome assembly v2.3. Read counts were assigned to genes and used to perform differential gene expression analysis using edgeR Bioconductor package v 3.38.4 (Robinson et al., 2010; McCarthy et al., 2012; Chen et al., 2016) on Rstudio (v 2022.07.1 Build 554) with R (v 4.2.1 (2022-06-23) -- "Funny-Looking Kid") using trimmed mean of M-values (TMM) for normalization (Robinson and Oshlack, 2010) and Negative Binomials Generalized linear models (GLMs) for statistical analysis. The p-values were adjusted using the Benjamini and Hochberg’s method to control the false discovery rate (FDR) (Benjamini and Hochberg, 1995). Enrichment analyses were performed using Mercator4 v2.0 (www.plabipd.de).

### Single-nucleotide polymorphism and Indel analyses

The detection of SNPs and Indels mutations in *N. caerulescens* accessions was performed using the “Basic Variant Detection” tool of CLC Genomics Workbench v.22.0.1 with *N. caerulescens* Cira genome assembly v2.3 as reference. RNA-seq reads from root samples were mapped to the annotated CDS using default parameters, except the length fraction = 0.8, the similarity fraction = 0.87 and ignoring non-specific matches. To call for mutations in the Basic Variant Detection tool, we used default parameters except for minimum coverage = 30, minimum count = 10, minimum central quality = 25, and minimum neighborhood quality = 20. The output was used to detect non-synonymous mutations using the Amino Acid Changes tool.

### Noccaea caerulescens grafting

The grafting of *N. caerulescens* shoots and rootstocks was performed essentially as previously described (Guimarães et al., 2009). Shoot scions and rootstocks were aligned on 1/2 MS plates containing 0.8% agar. The cotyledons of shoot scions were removed to maintain the alignment of the grafted tissues. Grafted seedlings were grown at 21°C with 16H light for 2 to 3 weeks. Successfully grafted plantlets were transferred to hydroponic culture and grown for 8 weeks in presence of 50 µM NiCl2 before elemental analysis of root and shoot samples.

### μXRF imaging of N. caerulescens roots

Seedlings of *Ncci* and *Nclc* were grown on modified Hoagland Agar plates supplemented with 50 μM NiCl_2_ for 8 weeks. Secondary roots were mounted in a homemade polycarbonate chamber (Mijovilovich et al., 2020), placed between a printer foil top window and a cellophane support and were kept hydrated with Hoagland nutrient solution excluding micronutrients. Optical images of the specimen were collected with an Optika microscope with a PRO8 digital camera (Ponteranica, Italy) prior to µXRF measurements. X-ray fluorescence spectra were collected with a customized M4 TORNADO µXRF imaging system (Bruker Nano GmbH, Berlin, Germany) equipped with two 50 mm^2^ active area silicon drift detectors as previously described (Mijovilovich et al., 2020). Briefly, an X-ray tube with Rh target with polycapillary optics with a 15 µm spot size at the manganese Kα line was operated at 50 kV and 600 µA. A filter composed of 100 µm of aluminum and 25 µm of titanium was used to flatten the Bremsstrahlung background. Stray counts were diminished by a customized shielding on the detector (Mijovilovich et al., 2020). The sample was measured with a step size of 4 µm (3 times oversampling) and a dwell time of 150 ms per pixel. XRF elemental images in 16 bits grayscale of micro and macro nutrients were obtained after spectral deconvolution and subtraction of the background in the BRUKER M4 software. Empirical references were made with a multi-element liquid standard contained in polyimide capillaries, as used in tomography experiments at synchrotrons (Mishra et al., 2013; Mijovilovich et al., 2019). These references have been quantified by ICP-MS. Elemental quantification was performed similarly to (Morina et al., 2021); all the computations were done with MATLAB R2019a at the computing and storage facilities MetaCentrum (Czech Republic). The 16-bit images in counts per second were multiplied by a “response factor” calibrated with the reference materials. The reference sample matched the thickness of the root. If a reference was not available for a certain root thickness, the value of the response factor was calculated from a regression curve. The oversampled pixels were binned (2x2), and a smoothing Gaussian filter (gamma 2) was applied. A false color map was applied using a customized not perceptually uniform color scheme (“HK7”) needed to visualize data with high variation among samples (Morina et al., 2021). Potassium images are semiquantitative (in counts after deconvolution).

### Molecular cloning of NcIRT1 promoter and coding sequences

All PCRs were performed using Phusion High-Fidelity DNA Polymerase (Thermo Fisher) and all constructions were confirmed by Sanger double stranded DNA sequencing or Oxford Nanopore sequencing. The sequence of the primers used in this study are given in (Supplementary Table S2). The promoter region of *NcIRT1* (g27500, 3.8 kb) was amplified by PCR from *Ncci* DNA using the primers proNcciIRT1-attB1 and proNcciIRT1-attB2. The PCR product was recombined into pDONR223 to produce pDON223-pro*NcIRT1*. The *NcIRT1* promoter was then re-amplified with primers proNcciIRT1-SalI and proNcciIRT1-XhoI. In parallel, the *GUSPlus* coding sequence was amplified from pCambia1035 using the primers GUSPlus-XhoI and GUSPlus-SalI. The two PCR fragments were digested with SalI and XhoI and cloned into pK7WG2D backbone (Karimi et al., 2002) opened with the SalI enzyme (thus removing the P35S and the Gateway cloning cassette), to generate pK7WG2D-pro*NcIRT1*-*GUSPlus*.

The predicted full-length coding sequence of *NcIRT1* was amplified by PCR from root cDNAs of *Ncfi* (*NcfiIRT1*) and *Nclc* (*NclcIRT1*) accessions with primers NcIRT1-attB1 and NcIRT1-attB2. The PCR products were recombined into pDONR207 and then pDR195-GTW (Oomen et al., 2009) to generate pDR195-*NcfiIRT1* and pDR195-*NclcIRT1* respectively.

The TagRFP coding sequence framed with 3xGA linkers was introduced in the first external loop of NcfiIRT1 at amino acid 41, as previously described for AtIRT1 (Dubeaux et al., 2018) using a PCR fusion-based approach. *NcIRT1* was amplified from pDR195-*NcfiIRT1* in 2 fragments with the pair of primers NcIRT1-attB1/NcfiIRT1-linkerGA-rev and NcfiIRT1-linkerGA-for/NcIRT1- attB2, respectively. The *TagRFP* coding sequence was amplified with primers TagRFP-linkerGA- for and TagRFP-linkerGA-rev. The 3 fragments were mixed and amplified with NcIRT1-attB1 and NcIRT1-attB2. The final fusion fragment was recombined into pDONR207 to generate pDONR207-*NcIRT1-TagRFP*. pDONR207-*NcIRT1-TagRFP* was then recombined with pK7WG2D to generate pK7WG2D-*NcIRT1-TagRFP*. For the complementation of the *A. thaliana irt1-1* mutant, entry vectors pDONRP4-P1R-proAtIRT1, pDONRP2R-P3-mock (Marquès-Bueno et al., 2016) and pDONR207-*NcfiIRT1* or pDONR207-*NcIRT1-TagRFP* were recombined with the pH7m34GW vector using multisite Gateway recombination to generate pH7m34GW- proAtIRT1-*NcfiIRT1* and pH7m34GW-proAtIRT1-*NcfiIRT1-TagRFP*.

### Functional analysis of NcIRT1 in Saccharomyces cerevisiae

The yeast *S. cerevisiae fet3fet4* mutant in the BY4741 strain was obtained by mating the *fet3* (Y06192) and *fet4* (Y16461) mutants (www.euroscarf.de). The double mutant was then complemented with the *HIS3* gene by homologous recombination to obtain the *fet3fet4H* strain used in this study (Mata; leu2Δ0; ura3Δ0; lys2Δ0; YMR058w::kanMX4; YMR319c::kanMX4). The *fet3fet4H* and BY4741H (García de la Torre et al., 2021) strains were transformed with pDR195, pDR195-NcfiIRT1, pDR195-NclcIRT1 and pDR195-AtIRT1 (provided by G. Vert laboratory) by the LiAc method and transformant selected on SD -Ura dropout agar medium. Complementation of *fet3fet4H* transformants was analyzed by drop assay on SD +Leu +Lys agar medium at pH 5.5 (10 mM MES), supplemented or not with 100 µM Fe-EDTA. Nickel sensitivity of the BY474H transformants was analyzed by drop assay on SD +Leu +Met agar medium at pH 5.5 (10 mM MES), supplemented or not with 500 µM NiCl2.

Metal accumulation was measured using three independent BY4741H transformant clones per construct. Saturated pre-cultures were diluted to OD600 = 0.4 in 50 mL SD +Leu +Met, 10 mM MES pH 5.5, 100 μM NiCl2, supplemented or not with 50 μM ZnSO4. Yeast cells were grown for 15 h at 28 °C with vigorous shaking. Yeast harvesting, washing, mineralization and elemental analyses were performed as previously described (González et al., 2024).

### Complementation of the Arabidopsis thaliana irt1-1 mutant

The *Arabidopsis thaliana irt1-1* mutant (Vert et al., 2002) was transformed with pH7m34GW, pH7m34GW-proAtIRT1-*NcfiIRT1* and pH7m34GW-proAtIRT1-*NcfiIRT1-TagRFP* by floral dipping using the *Agrobacterium tumefaciens* AGL0 strain (Clough and Bent, 1998). Transgenic T1 lines were selected on 1/2 MS agar medium supplemented with 25 μg.ml^-1^ hygromycin B (Duchefa Biochemie #H0192). T2 plants resistant to hygromycin B, together with WT (Ws) and *irt1-1* plants were grown in peat pots without addition of iron in greenhouse.

### Generation of Noccaea caerulescens transgenic lines

Five-day-old *Ncci* plantlets were transformed as previously described (Lin et al., 2016; García de la Torre et al., 2021) with the *Rhizobium rhizogenes* strain Arqua1 carrying the pK7WG2D- *proNcIRT1*-*GUSPlus* construct. Chimeric plantlets were grown on selection medium containing 50 μg.ml^-1^ kanamycin, and transgenic roots expressing GFP were selected from 2 to 6 weeks of culture using a Nikon AZ100 Fluorescence Macroscope, before analysis of GUS activity.

*Nclc* and *Ncci* plantlets were transformed using the same protocol with pK7WG2D and pK7WG2D-*NcIRT1-TagRFP* to express *NcIRT1-TagRFP* under the control of the 35S promoter. Chimeric plantlets were grown on selection medium containing 50 μg.ml^-1^ kanamycin, and roots were selected for GFP and TagRFP fluorescence. Selected *N. caerulescens* chimeric plantlets were transferred to hydroponic culture using a Hoagland nutritive solution containing 50 μM NiCl2 and grown in a climatic chamber (21°C, 16h light, 60% RH). Root systems were checked weekly for fluorescent signal to ensure the maintenance of the TagRFP and/or GFP expression. After 4 weeks, leaves and roots of each chimeric lines were harvested for elemental analysis and RT-qPCR analyses.

### Analysis of GUS activity

*Ncci* roots transformed with pK7WG2D-pro*NcIRT1*-*GUSPlus* and expressing GFP were stained for GUS activity essentially as previously described (Fiorucci et al., 2022). Roots were fixed with ice-cold 90% acetone under vacuum for 15 min and then washed twice with 50 mM sodium phosphate buffer pH 7.2. Roots were then vacuum infiltrated three times for 5 min with modified a 5-bromo-4-chloro-3-indolyl-β-glucuronide (X-Gluc) staining solution [50 mM NaPO4 pH 7.2, 0.1% Triton X-100, 0.5 mM K3[Fe(CN)6], 0.5 mM K4[Fe(CN)6], 2mM X-Gluc] on ice in the dark. GUS activity was revealed by incubation at 37°C for 5 to 10 min. Root tissues were cleared by incubation in 70% ethanol at 4 °C, and both GFP fluorescence and GUS staining were imaged using a Nikon AZ100 Fluorescence Macroscope.

### Cellular localization of NcIRT1

*Nclc* chimeric plantlets transformed with pK7WG2D-*NcIRT1-TagRFP* and expressing TagRFP were transferred to Hoagland agar plates containing 50 μg.ml^-1^ kanamycin, supplemented or not with 50 μM NiCl2, and grown for an additional two weeks. Localization of NcIRT1-TagRFP was analyzed at the IMAGERIE-Gif platform (www.i2bc.paris-saclay.fr/bioimaging/) using a Leica SP8X inverted confocal microscope with laser excitation at 555 nm and collection of emitted light at 570-620 nm.

### Elemental analyses of N. caerulescens samples

*N. caerulescens* leaf and root samples from hydroponic cultures were washed twice with 10 mM CaCl2/2mM MES-KOH pH 5.7 and rinsed twice with ice-cold ultrapure water. Samples were dried at 65 °C for 48 h and then weighed to determine the dry mass. Dried samples were mineralized in 2 mL of 65 % HNO3 for 4 h at 120 °C, and for an additional 4 h after the addition of 1 mL of 30 % H2O2. Mineralized samples were analyzed on an Agilent 4200 MP-AES (Agilent Technologies) and metal concentration was calculated by comparison with a metal standard solution.

### Analysis of IRT1 transgene expression in hairy root lines

Roots of individual *Nclc* plantlets transformed with pK7WG2D-*NcIRT1-TagRFP* and grown in hydroponic culture in presence of nickel were frozen in liquid nitrogen. Frozen roots were grinded and total RNA was extracted using the RNeasy Plant Mini Kit with DNase I treatment according to manufacturer instructions (Qiagen). cDNAs were synthetized by random priming from 300 ng RNA using the SuperScript IV First-Strand Synthesis System (Thermo Fisher Scientific). The *IRT1-TagRFP* transgene was specifically amplified with primers NcIRT1-TagRFP-for and NcIRT1-TagRFP-rev. The 6-phosphogluconate decarboxylating 3 gene (*Nc6PGDC,* g11049 in *Ncci* genome v2.3) used as reference gene was amplified with primers Nc6PGDC-for and Nc6PGDC-rev (García de la Torre et al., 2021). Quantitative PCR analyses were performed on a LightCycler 96 using LightCycler® 480 SYBR Green I Master (#04707516001, Roche) and gene expression was quantified with the 2^ΔΔCt^ method (Livak and Schmittgen, 2001).

### Statistical analyses

Statistical and Pearson correlation analyses were performed using Prism software v. 7.05 with p-value < 0.01.

## Acknowledgments

We deeply thank Dr. Ana Mijovilovich (Institute of Plant Molecular Biology, Laboratory of Plant Biophysics & Biochemistry, České Budějovice) for her invaluable contribution and expertise with µXRF experiments. Bojan Vujić is acknowledged for 3D printing of chambers for in vivo µXRF measurements and sample preparation tools for μXRF. We also thank Sophia Rafasse for generating the yeast *fet3fet4* double mutant, Dr Claude Grison for providing the seeds of the *N. caerulescens* Bergenbach accession, and the laboratory of Grégory Vert for providing us the pDONRP4-P1R-proAtIRT1, pH7m34GW and pDR195-AtIRT1 constructs. We are very gratefull to the Dirección Xeral de Patrimonio Natural - Xunta de Galicia (Spain) for the authorization to collect seeds of *N. caerulescens* Puente Basadre and Cira accessions (# EB-005/2023), and the prefecture of Aveyron (France) for the access to the seeds of the Firmiensis accession (#12-2014- 06).

## Author contributions

**Conceptualization:** CB, VSG, SM

**Performed experiments:** CB, VSG, RCA, HK, CR, CI, CV, ALA, WL, JY

**Analyzed data:** CB, VSG, RCA, HK, CK, JH, MGMA, SM

**Provide resources:** MGMA, CQS

**Funding acquisition:** HK, MGMA, ST, SM

### Writing, original draft: CB, SM

**Writing, review and editing:** CB, VSG, RCA, HK, CV, CK, MGMA, CQS, ST, SM

## Data availability

The Genome assembly of *Noccaea caerulescens* Cira (v2.3) is available at NCBI under Bioproject PRJNA1096268, and the corresponding raw sequencing data available as SRA Biosample SAMN40749631. The annotation files for the *N. caerulescens* Cira genome assembly v2.3 are available at Zenodo (doi: 10.5281/zenodo.11371499). *N. caerulescens* genome resequencing data for Prayon, Plombières and La Calamine accessions are available at NCBI SRA under accessions SRAXXXXXXXX, SRAXXXXXXXX and SRAXXXXXXXX.

Transcriptomic data for *N. caerulescens* accessions and *Microthlaspi perforliatum* are available under NCBI Bioproject PRJNA657163.

