## Supplemental Figures for "Gain and loss of gene function shaped the nickel hyperaccumulation trait in *Noccaea caerulescens*"

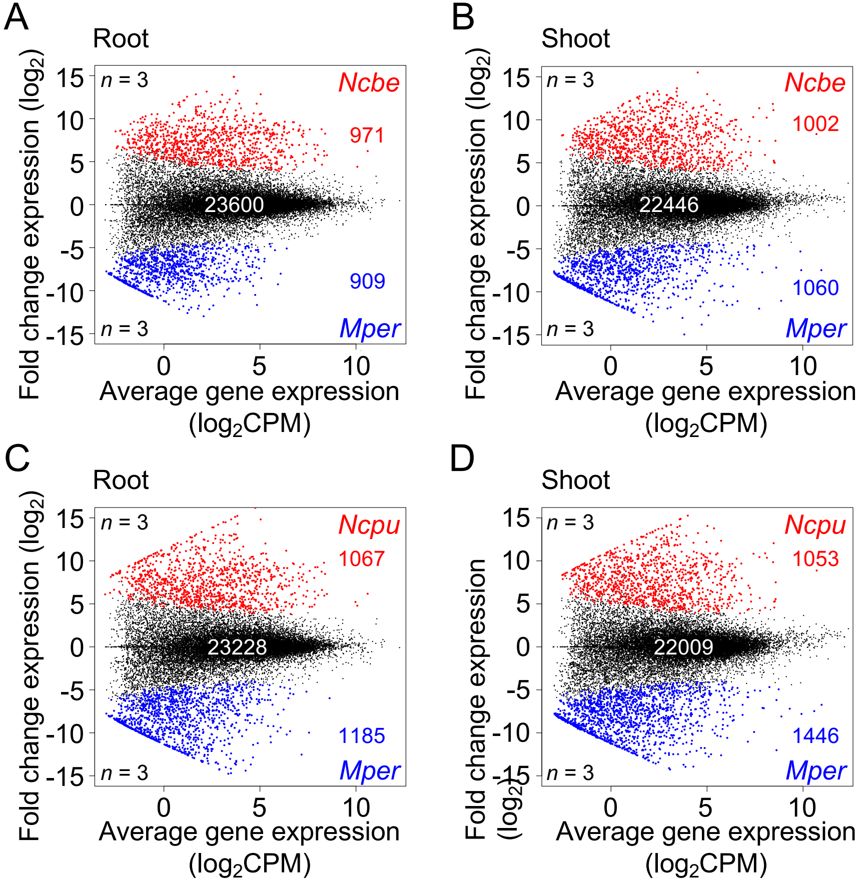


**Supplementary Figure S1.** Identification of differentially expressed (DE) genes between *Noccaea caerulescens* and *Microthlaspi perfoliatum.* A) MA plot representing the interspecific comparative transcriptomics analysis of gene expressed in roots of *N. caerulescens* Bergenbach (*Ncbe*) and *M. perfoliatum* (*Mper*). B) MA plot representing the interspecific comparative transcriptomics analysis of gene expressed in shoots of *Ncbe* and *Mper*. C) MA plot representing the interspecific comparative transcriptomics analysis of gene expressed in roots of *N. caerulescens* Puente basadre (*Ncpu*) and *M. perfoliatum* (*Mper*). D) MA plot representing the interspecific comparative transcriptomics analysis of gene expressed in shoots of *Ncpu* and *Mper*. In all comparisons, the number of DE genes ($|log_2FC| \geq 3.32$, FDR adjusted p-value $< 0.05$, $n = 3$ biological replicates), more expressed in *N. caerulescens* accessions (red dots) and in *Mper* (blue dots) are indicated.

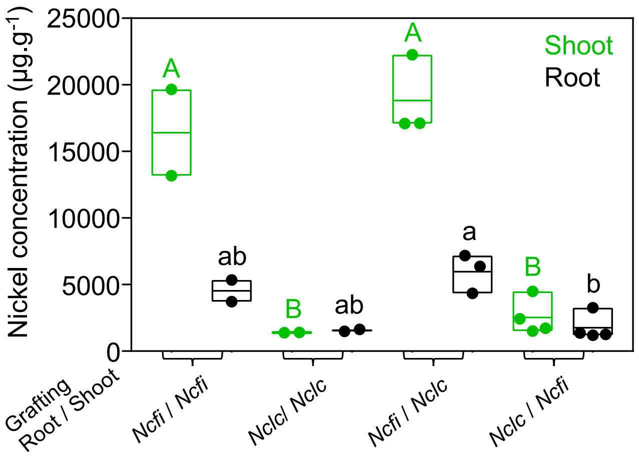


**Supplementary Figure S2.** Origin of the difference in metal accumulation between *Noccaea caerulescens* Firmiensis (*Ncfi*) and La calamine (*Nclc*) accessions. Nickel accumulation in shoots (green) and roots (black) of plants derived from the self or reciprocal grafting of rootstock and shoot explants of *Ncfi* and *Nclc* accessions. Individual and mean metal concentration values are presented. Significant differences between the means are indicated by different letters for shoots and roots ($n$ = 2 - 4 biological replicates, one-way ANOVA with Tukey's multiple comparison test after logarithmic transformation of the data, p-value < 0.01).

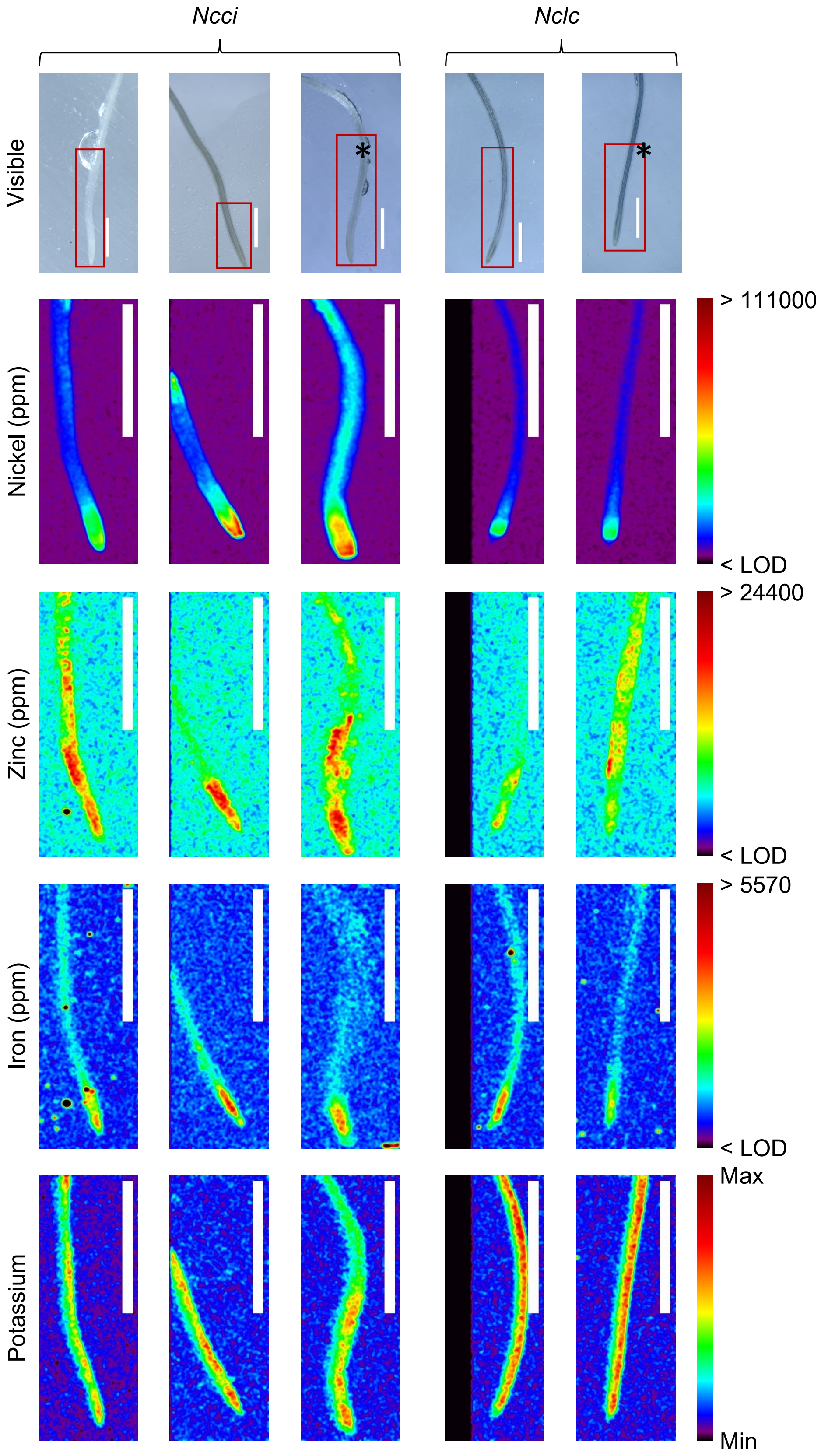


**Supplementary Figure S3.** Metal distribution in roots of *Noccaea caerulescens* Cira *(Ncci)* and La calamine *(Nclc)* accessions. µXRF imaging of *Ncci* and *Nclc* roots for analysis of nickel, zinc, iron and potassium semiquantitative distribution. The red square on the visible images (top lane) represents the area analyzed by µXRF for the quantification of elements. When indicated (LUT on the right), the XRF images have been calibrated for quantification (ppm) with XRF standard. * denotes the replicates that are presented in Fig. 4C. Scale bar : 1 mm.

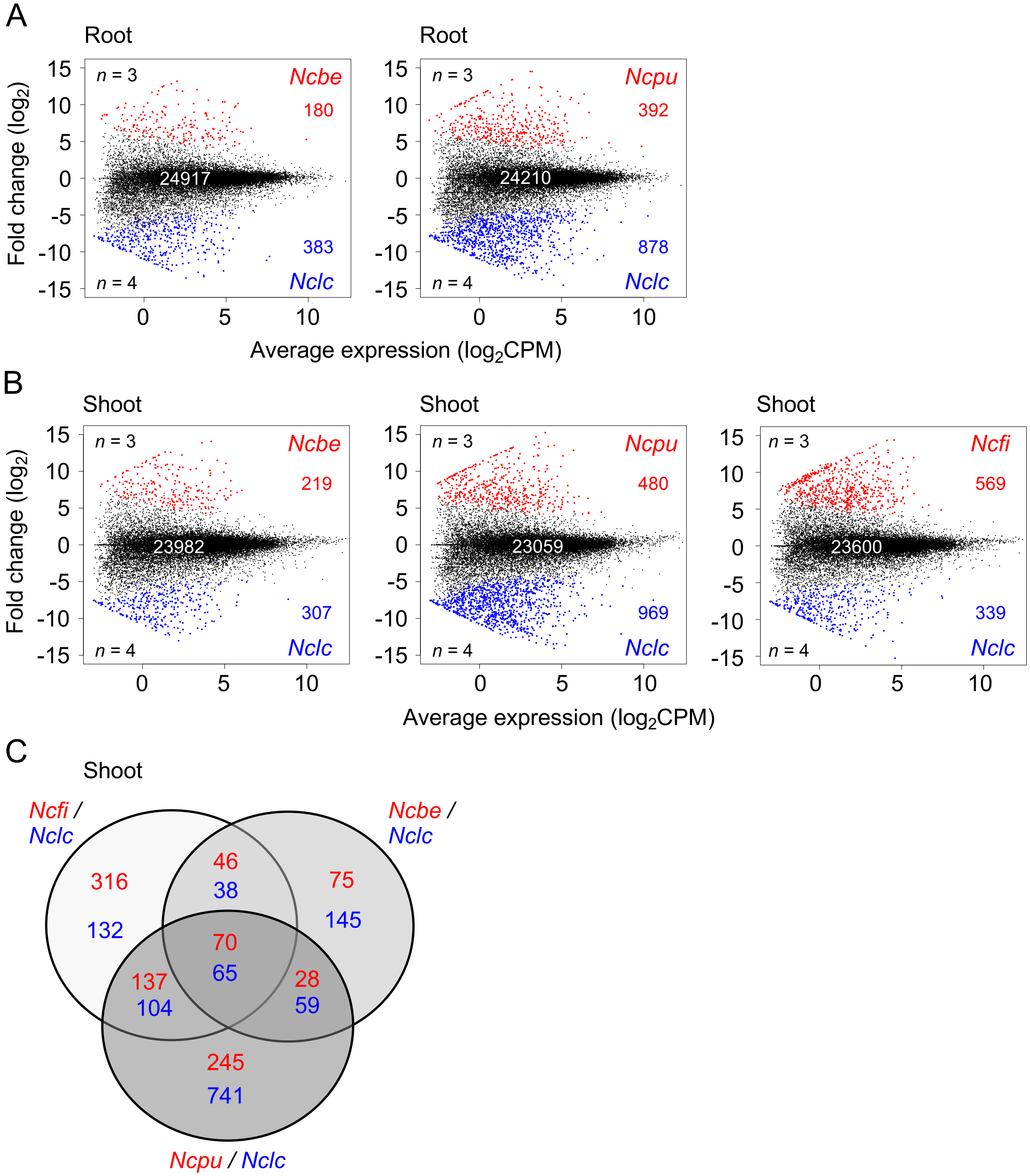


**Supplementary Figure S4.** Intraspecific comparative transcriptomics of *Noccaea caerulescens* in roots and shoots. A) MA plots representing the intraspecific comparative transcriptomics analysis of genes expressed in roots of *N. caerulescens* Bergenbach (*Ncbe*) and Puente basadre (*Ncpu*) with La calamine (*Nclc*). The number of differentially expressed (DE) genes ($|\log_2 FC| \geq 3.32$, FDR-adjusted p-value $< 0.05$, $n$ = 3-4 biological replicates), more expressed in *Ncbe* or *Ncpu* (red dots) and in *Nclc* (blue dots) are indicated. B) MA plots representing the intraspecific comparative transcriptomics analysis of genes expressed in shoots of *N. caerulescens* Ncfi, *Ncbe,* and *Ncpu* with *Nclc.* The number of DE genes ($|\log_2 FC| \geq 3.32$, FDR-adjusted p-value $< 0.05$, $n$ = 3-4 biological replicates), more expressed in Ncfi, *Ncbe* or *Ncpu* (red dots) and in *Nclc* (blue dots) are indicated. C) Venn diagram displaying the number of DE genes in shoots between the nickel hyperaccumulating accessions *Ncfi*, *Ncbe*, *Ncpu* (red) and *Nclc* (blue)

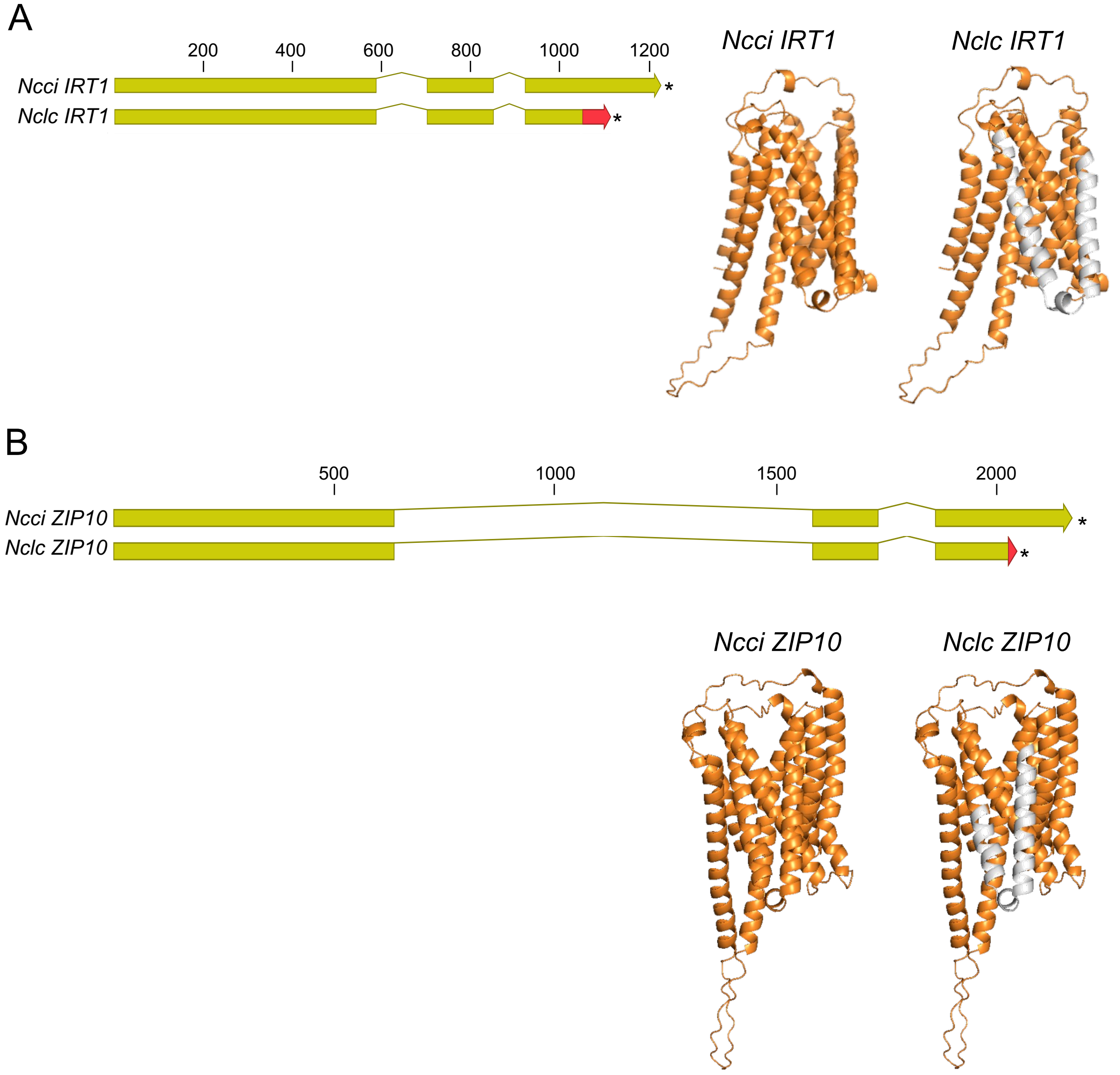


**Supplementary Figure S5.** Mutations affecting the *NcIRT1* and *NcZIP10* coding sequences in *N. caerulescens* La calamine (*Nclc*). A) Structure of *NcIRT1* (g27500) and B) *NcZIP10* (g22398) coding sequences and consequences of the frameshift mutations identified in *Nclc* compared to the Cira (*Ncci*) reference (see Table 3). The red color in the CDS indicates changes in amino acids, and the asterisks the early stop codon. The consequences of these mutations are presented on the structural model of IRT1 and ZIP10 obtained by AlfaFold (https://alphafold.ebi.ac.uk). The grey color in the protein models indicates the predicted missing domains.

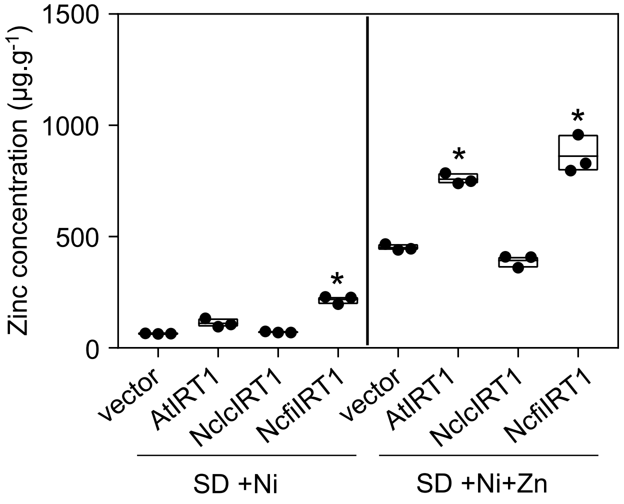


**Supplementary Figure S6.** *NcIRT1* expression increases zinc accumulation in yeast. NcfiIRT1, as well as NclcIRT1 and AtIRT1, were expressed in BY4741H yeast cells and zinc accumulation was measured in transformants grown in liquid SD medium containing 100 µM $NiCl_2$ (SD +Ni) or in the same medium supplemented with 50 µM $ZnSO_4$ (SD +Ni+Zn) as in Fig. 6D. Yeast cells transformed with the empty pDR195 were used as control. The data represent individual measurements as well as the mean, minimum, and maximum values. Asterisks indicate significant differences compared to the control (vector) mean (Welch's t-test after logarithmic transformation of the data, p-value < 0.01, n=3).

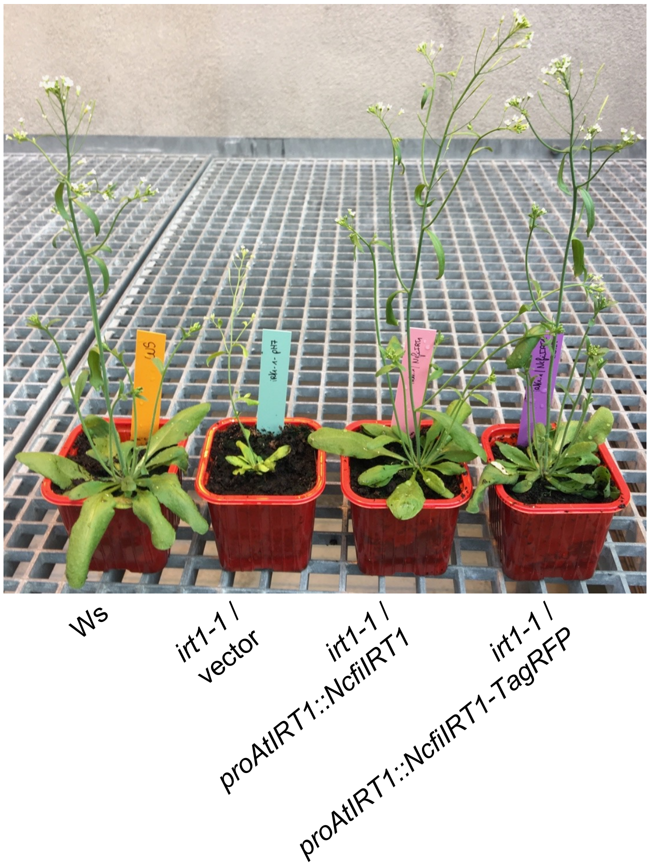


**Supplemental Figure S7.** Complementation of the *Arabidopsis thaliana irt1-1* mutant with NcfiIRT1. *N. caerulescens* IRT1, cloned from the Firmiensis accession (NcfiIRT1), as well as the TagRFP tagged version (NcfiIRT1-TagRFP) were expressed in the *A. thaliana irt1-1* mutant under the control of the *A. thaliana* IRT1 promoter (proAtIRT1) using the pH7m34GW vector. T2 transformants resistant to hygromycin B, together with WT plants (Ws accession) were grown in peat pots in greenhouse without addition of iron.
